# Where are they now? Academic and career trajectories of national laboratory STEM internship alumni from community colleges, compared to those from universities

**DOI:** 10.1101/2025.03.24.645145

**Authors:** Laleh E. Coté, Astrid N. Zamora, Julio Jaramillo Salcido, Seth Van Doren, Aparna Manocha, Gabriel Otero Munoz, Esther W. Law, Anne M. Baranger

## Abstract

It is well-established that participation in technical or research experiences can support university student retention in science, technology, engineering, and mathematics (STEM). However, there is very little published research on the availability and impact of opportunities for community college students, particularly those provided by Department of Energy (DOE) national laboratories. To address this gap, we collected data from students who participated in two DOE sponsored programs spanning from 2009 to 2016: the Community College Internship (CCI) and Science Undergraduate Laboratory Internship (SULI) at Lawrence Berkeley National Laboratory (LBNL). Among CCI alumni, 90% earned a STEM bachelor’s degree and 88% are on a STEM career pathway. For SULI alumni, 91% earned a STEM bachelor’s degree and 71% are on a STEM career pathway. Overall, 80% of CCI alumni and 56% of SULI alumni have entered the STEM workforce, 5% of CCI alumni and 11% of SULI alumni are in the health workforce, and 6% of CCI alumni and 13% of SULI alumni are in the non-STEM workforce. Our findings indicate that community college students who participate in STEM professional development activities (such as the CCI program) are likely to complete their academic degrees and pursue STEM careers at rates comparable to those of university students. This investment in providing internships at LBNL for community college students has effectively supported their entry into STEM careers and their desire in pursuing work within the DOE complex in the years following their participation in these programs.

## Introduction

To support scientific innovation, technological advancement, and the national economy, the United States (U.S.) will need to continue its investment and attention to the development of the science, technology, engineering, and mathematics (STEM) workforce (NASEM, 2023; NLDC, 2024). For the past decade, there has been a strong focus on increasing the number of U.S. trainees in STEM fields, which has been featured prominently in various federal calls to action (America COMPETES Act of 2022, 2022; PCAST, 2012, 2021). Due to concerns about a shortage of STEM-trained professionals and the collaborative advantages of teams representing different career stages and backgrounds, in recent years there has also been a growing focus on the “aging workforce” in STEM (Blau & Weinberg, 2017; Byrd & Scott, 2018; Conway & Monks, 2017; Lim et al., 2017; Sinnott et al., 2021; Smith-Doerr et al., 2017; White et al., 2018; Zerhouni et al., 2016). Federal agencies such as the U.S. Department of Defense (DOD), U.S. Department of Energy (DOE), National Aeronautics and Space Administration (NASA), National Institute of Standards and Technology (NIST), National Science Foundation (NSF), and U.S. Department of Veterans Affairs (VA) are important contributors to major scientific and technological discoveries, raising awareness about the importance of the STEM workforce, and providing opportunities to explore STEM careers for people at various education levels and career stages.

Established in 1977, the DOE is the largest federal sponsor of physical sciences research, leads the nation’s strategy in clean energy technologies, and carries out much of its environmental science, nuclear security, and energy work at its 17 national laboratories (DOE, n.d.; EWAB, 2024). With bipartisan support, the “concentration of scientific expertise and infrastructure” at the DOE national laboratories is viewed as integral to its scientific and technological contributions (Office of Congressman Bill Foster, 2018; U.S. Senate, 2023; Widener, 2013). In this paper, we focus on addressing the needs of the DOE national laboratory system, which anticipates staffing challenges and loss of institutional knowledge due to the impending retirement of a large proportion of its workforce (DOE, 2017; Energy Workforce Opportunities and Challenges, 2019; National Research Council, 2015). As a way to address this issue, the DOE supports internships and professional development opportunities for students at DOE national laboratories and facilities throughout the U.S. Ultimately, the DOE hopes to retain some of these students as future employees (DOE, 2017, 2022; DOE SC, 2020; McDowell et al., 2024). The DOE Office of Science (SC) Office of Workforce Development for Teachers and Scientists (WDTS) hosts a suite of national programs for undergraduates, post-baccalaureates, graduate students, and faculty members.

For example, the **Science Undergraduate Laboratory Internship (SULI)** program provides STEM internship opportunities to undergraduates from community colleges^1^ and baccalaureate granting institutions^2^, but serves more individuals from the latter. The **Community College Internship (CCI)** program is similar to the SULI program in many respects, though it is only open to undergraduates from community colleges who have completed STEM coursework. As the country’s primary mechanism for workforce training, the U.S. community college system has been identified as critical to the education and training of the future STEM workforce (CCRC, 2021; Jacobs & Worth, 2019; PCAST, 2021). Community colleges are a valuable resource in the higher education system, as they are located in every region of the U.S., support local economies, and provide educational opportunities to people from all backgrounds (e.g., AACC, 2024; Camardelle et al., 2022; D’Amico et al., 2019; Jacobs, & Worth, 2019; NCES, 2022). For example, military veterans choose community colleges over other higher education institutions, in part because of the flexibility of course schedules and availability of student services (Jones, 2017). Similarly, many student parents with caregiving and financial obligations, find community colleges to be a favorable option (Cruse et al., 2019; Reed et al., 2021). Thus, the current study is focused on CCI, and uses SULI as a point of comparison.

Previous work by Foltz and colleagues (2011) recommends the evaluation of DOE program outcomes related to “educational advancement and workforce development.” This work also highlighted the importance of collecting data on “long-term outcomes,” and maintaining communication and engagement with program alumni. Similarly, more recent reports from the DOE have called for the collection of quantitative data to assess the impact of its “educational and community outreach efforts,” engage in longitudinal tracking of program participants, determine program participant entry into the DOE workforce, and determine the success of programs in promoting careers in STEM (DOE, 2022; DOE SC, 2020). In the current study, we address these recommendations by examining academic and career activities in the years following undergraduates’ participation in two DOE internship programs.

As we will describe in the literature review, STEM technical and research internships have the potential to support undergraduate learning, academic success, and retention in STEM (e.g., Haeger et al., 2024). These benefits have been well-documented for undergraduates at baccalaureate granting institutions, especially for students from groups historically excluded from STEM fields. However, there are relatively few studies that include details about how professional development opportunities impact community college students’ academic achievement and career pathways (e.g., Bulman & Fairlie, 2022; Crisp et al., 2016; Monaghan & Sommers, 2022; Taylor & Jain, 2017). Given the importance of community colleges in developing the future STEM workforce and the limited research on community college students’ academic and career pathways in STEM fields, our central focus was to examine the academic and career activities of CCI participants – community college STEM majors – in the years following their completion of the program. To better understand this unique population, we collected data on academic and career activities and interest in working at a DOE national laboratory, from both CCI alumni (our study population) and SULI alumni (our comparison group). We focused on the following research questions:

***Research Question 1:*** *What are the academic and career trajectories of Community College Internship (CCI) alumni in the years following their participation in the program at Lawrence Berkeley National Laboratory (LBNL)?*
***Research Question 2:*** *How do the academic and career trajectories of CCI alumni compare with those of a similar program, the Science Undergraduate Laboratory Internship (SULI) program at LBNL?*

Although there are relatively few studies on community college academic and career activities, even *fewer* compare the activities of community college or transfer students with those of students who began their studies at baccalaureate granting institutions (e.g., Xu et al., 2018). Thus, we examined the percentage of CCI alumni who graduated with degrees, entered graduate programs, and joined the workforce. Our previous work on CCI alumni highlighted the unique learning environments of DOE national laboratories; therefore, we also examined the percentage of CCI alumni currently working at a DOE national laboratory or facility, or who are interested in doing so in the future (Coté et al., 2025).

## Literature review

### Technical/research experiences provide academic and career benefits to undergraduates

Undergraduates who believe their participation in activities has contributed to learning, personal growth, and the development of positive relationships with faculty or professionals in their field are more likely to have a favorable view of their undergraduate education and career prospects (Andrade et al., 2022; Johnson & LaBelle, 2022; Torpey-Saboe & Clayton, 2023). Many students are motivated to participate in technical/research experiences to gain new skills or knowledge or because they are curious about graduate schools and/or careers in a particular field (Coté et al., 2025; Craney et al., 2011; Nolan et al., 2020; Trott et al., 2020). Many scholars agree that participation in a mentored internship or research experience reduces degree completion time and increases academic achievement, STEM degree completion rates, interest in pursuing a STEM graduate degree, and persistence in STEM, particularly for individuals from groups who have been historically excluded from STEM, which can be categorized by factors such as race, ethnicity, gender, and socioeconomic status (Chemers et al., 2011; Eagan Jr et al., 2013; Haeger et al., 2024; Hernandez et al., 2018; Hewlett, 2016; Jelks & Crain, 2020; Nerio et al., 2019; Prunuske et al., 2016; Romero et al., 2023). Aligned with career success over time, students benefit from completing technical/research experiences when they develop interpersonal relationships with members of the STEM community, including graduate students and professionals (Romero et al., 2023; Trott et al., 2020; Zydney et al., 2002). This professional network can be leveraged over the course of their career, and lead to additional opportunities.

### There is little scholarship about technical/research experiences for community college students

Although nearly half of all students with STEM degrees in the U.S. attend a community college at some point during their educational careers, there is limited research focused on internships and/or research experiences for community college students and/or students who transferred from a community college to a baccalaureate granting institution (Creech et al., 2022; Krim et al., 2019; Lucero et al., 2021; Linn et al., 2015; Mooney & Foley, 2011; NASEM, 2017b; NSF, 2011; Tuthill & Berestecky, 2017; Tsapogas, 2004). Community colleges enroll disproportionately high numbers of Black, Hispanic/Latinx, Native American, and disabled students, but these groups have very low rates of retention, completion of a certificate or degree, or transfer to a baccalaureate granting institution (Camardelle et al., 2022; Genthe & Harrington, 2022; Madaus et al., 2021; The Campaign for College Opportunity, 2021; Torpey-Saboe & Clayton, 2023). Thus, participation in technical/research internships could provide community college students from all backgrounds with experiences and resources to support their academic and career success.

### Community colleges play an important role in STEM workforce development

In the U.S., community colleges provide opportunities to develop skills or certifications needed to enter the workforce through middle-skill jobs or transfer to a baccalaureate granting institution. However, these are not the only motivations for enrollment (Belfield & Brock, 2023; Cohen & Kelly, 2019). Projected to increase in number and availability, middle-skill jobs are those that require certification beyond a high school diploma but less than a bachelor’s degree and are an important component of the future STEM workforce (Carnevale et al., 2018; NASEM, 2017a; Paz-Soldan et al., 2025). Middle-skill jobs include electricians, hydrologic technicians, industrial machinery mechanics, nuclear reactor operators, solar thermal installers, telecommunications engineering specialists, and wind facilities managers (O*NET, 2024). Community college students who transfer to baccalaureate granting institutions to obtain bachelor’s degrees can enter the STEM workforce in jobs such as agricultural engineers, biodiesel division managers, chemical engineers, data scientists, field engineers, microbiologists, supply chain managers, and system programmers (O*NET, 2024). The DOE has identified a critical need to develop energy and domestic manufacturing workforces, which will require engagement and partnerships with community colleges across the U.S. (EWAB, 2024). Beyond the DOE complex, numerous reports examining the untapped potential of the U.S. education system to support the development of the STEM workforce, highlighting the need to leverage community colleges as part of this overall strategy (e.g., McDowell et al., 2024; Paz-Soldan et al., 2025). In fact, 22% of U.S. workers who received their education from the community college system are part of the STEM workforce, illustrating the potential value of gaining a deeper understanding of this population (Belfield & Brock, 2023).

### Programs hosted at national laboratories are unique, yet understudied

In collaboration with more than 450 academic institutions across the U.S. and Canada, DOE national laboratories invest over $500 million each year to support students, recent graduates, postdoctoral fellows, and faculty members through sponsored research experiences. However, these institutions are almost entirely absent from the extensive body of educational research literature on such programs (DOE, 2017; Krim et al., 2019). To our knowledge, only one project – a doctoral capstone – has examined how DOE internships compare to other federally funded programs and evaluated their outcomes (Foltz et al., 2011). Conducted in partnership with DOE Office of Science (SC) Office of Workforce Development for Teachers and Scientists (WDTS) staff members, this project collected survey data from alumni of the Science Undergraduate Laboratory Internship (SULI) and the joint NSF and National Institute of Standards and Technology (NIST) Summer Undergraduate Research Fellowship (SURF) programs.

As compared to the typical internship or research experience highlighted in the literature, programs at national laboratories are unique in that a) they are not situated at a college or university, b) individuals who participate as mentors typically are scientists, engineers, technical staff, and postdoctoral researchers rather than faculty and graduate students, c) some projects give students access to large-scale interdisciplinary research facilities beyond the capability of the typical faculty-led research group housed at a college or university. Similar to Research Experiences for Undergraduates (REU) programs funded by the NSF, internships at national laboratories collectively engage thousands of participants each year from colleges and universities across the U.S., in a variety of disciplines (Foltz et al., 2011). This study is one of few academic publications that include *both* a) data from DOE national laboratory program participants and b) documentation of human subjects committee review/approval at the host institution (e.g., Coté et al., 2025; Fong et al., 2025; Ivory et al., 2023). When this information is included in a study, we anticipate new opportunities to increase the visibility of programs hosted by DOE national laboratories, and create more awareness about their impacts on STEM education in the U.S. Publishing findings about programs and other activities in support of the STEM workforce at DOE national laboratories in peer-reviewed journals and conference proceedings will enable scholars from other institutions to *access, contribute* to, and *engage with* this information.

### Tracking participants’ long-term academic and career activities illustrates program impacts

Numerous studies on STEM research experiences have been published in recent decades. However, they primarily focus on the experiences and outcomes of students enrolled at baccalaureate granting institutions (Krim et al., 2019; Linn et al., 2015; NASEM, 2017b). Many studies designed to measure the impacts of research experiences on student success and persistence in STEM are merely descriptive or do not track student success beyond the acquisition of an undergraduate-level degree (Estrada et al., 2018; Krim et al., 2019). In fact, there is little tracking of STEM majors’ long-term career outcomes in existing literature, as few studies report on the long-term impacts beyond 4 years after a research experience (Dou et al., 2021; Trott et al., 2020). Commonly, studies about the impacts of “science training programs” rely on short-term outcomes such as levels of motivation, interest, and intention to stay in STEM, which are important, but would be more powerful when coupled with longitudinal data about academic and career activities (Estrada et al., 2021). Thus, in this study, we have examined data between 5 and 12 years after students’ participation in the CCI and SULI programs at LBNL, and have centered our efforts on understanding community college students’ academic achievement and retention in STEM.

## Description of Programs

In support of the DOE mission – to ensure America’s security and prosperity by addressing its energy, environmental, and nuclear challenges through transformative science and technology solutions – DOE SC WDTS collaborates with national laboratories and other facilities to sponsor and manage a suite of internship programs. Two of these, the **Community College Internship (CCI)** and **Science Undergraduate Laboratory Internship (SULI)** programs, are highlighted in this study. These programs are designed to encourage students and recent graduates to enter STEM careers through immersive technical and research experiences at DOE laboratories at a host laboratory/facility in the spring, summer, or fall terms. Both CCI and SULI support the development of the DOE workforce, and national laboratories engage in outreach efforts to increase the likelihood that students and recent graduates from a variety of backgrounds apply to participate (Hampton-Marcell et al., 2023; DOE SC, 2020). Eligible applicants are those who are 18 years or older at the start of the program. For those applicants who have not yet completed their undergraduate studies, they must be enrolled full-time at an undergraduate institution, have a minimum Grade Point Average of 3.0 on a 4.0 scale, and have completed at least six credit hours of STEM coursework and 12 hours of coursework.

These programs are hosted by national laboratories across the U.S., but this study focuses on one laboratory in particular, Lawrence Berkeley National Laboratory (LBNL). At LBNL, Workforce Development & Education serves as the host for several internships, including CCI and SULI. Participating undergraduates and recent graduates are referred to as “interns.” Each intern is placed with a “Mentor Group,” which consists of those individuals responsible for the daily supervision, teaching, and mentoring of interns and their colleagues. Altogether the team may consist of permanent staff, temporary staff, postdoctoral scholars, and/or graduate students, though the structure of these teams varies widely. In support of intern development from novice to expert, these Mentor Groups engage in both teaching (e.g., technical skills training) and mentoring (e.g., sponsorship, conversations about STEM careers) over the course of the internship term (Helix et al., 2022; Zalaquett & Lopez, 2006).

Interns attend orientation, meetings, and training sessions throughout the term. An internship coordinator facilitates check-in meetings to discuss interns’ experiences, address any issues, and connect them with resources (e.g., online training courses, health services, computer equipment, information about events) at LBNL. They have opportunities to attend optional enrichment activities, such as tours and seminars. Interns also attend workshops during which they learn about effective presentation techniques. Throughout the program, interns complete a technical report or research paper and poster and present at a poster session during the final week of the program.

Interns receive a stipend and housing supplement and are reimbursed for travel costs to and from LBNL. They are eligible to receive these funds by submitting any required “onboarding” paperwork, dedicating 40 hours a week to the internship for the duration of the program, participating in mandatory events, and completing written deliverables. Prior to 2013, interns received $500 per week, which included both stipend and housing. In 2013, interns received a stipend of $16.25 per hour ($650 per week) and $150 per week for housing costs. In 2016, interns received a stipend of $16.25 per hour ($650 per week) and $300 per week for housing. For comparison, the minimum wage in California was $8.00 per hour in 2013, $9.00 per hour in 2014, and $10.00 per hour in 2016 (State of California Department of Industrial Relations, 2023). Providing financial support is important, because students who need to work during their undergraduate studies to fund their education are *less* likely to participate in unpaid professional development opportunities (Coté, 2023; Pierszalowski et al., 2021; Pratt et al., 2019; Drake et al., 2019). Additionally, paid opportunities can support students’ self-esteem, self-efficacy, feelings of being valued by the sponsor, interest in participation, and likelihood of staying in school (Coté et al., 2025; Minasian, 2019; Pratt et al., 2019; Romero et al., 2023).

## Methods

### Author expertise and background

As an undergraduate biology major, L.E.C.^3^ attended multiple community colleges in California and completed the CCI program in 2007 and the SULI program in 2008 and 2009. She worked with the same Mentor Group at LBNL during these internships and for an additional 2 years as a Research Assistant. L.E.C. has 14 years of experience implementing and managing various internship programs hosted by Workforce Development & Education at LBNL, including CCI and SULI. A.M.B. has 28 years of experience working with and studying STEM undergraduate research experiences. L.E.C. and A.M.B. have worked together for 9 years, studying this topic, generating curricula for training workshops about teaching and mentoring undergraduate researchers, and working to improve undergraduate-level courses in biology and chemistry. As student assistants or staff, E.W.L., J.J.S., and S.V.D. worked at LBNL with internship programs hosted by Workforce Development & Education at LBNL. A.M., A.N.Z., and G.O.M. contributed to this project as undergraduate and graduate research assistants through partnerships with the University of California, Berkeley. The authors of this work include women and men, whose backgrounds include Asian, Black, Hispanic, Latinx, Middle Eastern, and/or White. Some authors identify as mixed race.

### Data collection

Our primary mode of communication was email. We sent out emails to all individuals for whom we had contact information and who had completed the program in 2009-2016. This recruitment method was not purposive, as the sample was not selected based on specific characteristics but rather included everyone for whom contact details were available. However, some contact information was missing or inactive (e.g., temporary email addresses from undergraduate institutions). For those individuals, we attempted to reach them through online social media platforms, such as Facebook and LinkedIn. We administered consent forms and surveys to potential study participants through Qualtrics. These methods resulted in data collection from 86 CCI alumni (out of a total of 93 individuals who completed CCI during 2009-2016: retention rate: 92.5%) and 90 SULI alumni (out of a total of 99 individuals who completed SULI during 2009-2016; retention rate: 90.9%). The data collection and follow-up communication with study participants occurred between 2018 and 2021. A.M., A.N.Z., E.W.L., G.O.M., J.J.S., L.E.C., and S.V.D. maintained and organized data throughout the project. As shown in **Figures S1** and **S2** in the Supplemental Materials, we asked CCI and SULI alumni to respond to survey questions about their professional activities across the DOE complex in the years following their internship at LBNL, and interest in working at a DOE national laboratory or facility. These methods were approved by the Institutional Review Board at LBNL (Protocol ID: Pro00023065) with the University of California, Berkeley, as the relying institution (Reliance Registry Study #2593). All contributing researchers have completed training in the responsible and ethical conduct of research administered through LBNL or the University of California, Berkeley.

### Study population and comparison group

The study included two groups of alumni. The first group is our **study population**, consisting of 93 individuals who completed the CCI program at LBNL between the Summer of 2009 and the Fall of 2016. These individuals were community college students at the time of their participation in CCI at LBNL, enrolled in institutions located across the U.S. in Alabama, Arizona, California, Florida, Illinois, Massachusetts, Nevada, New York, and Washington. We collected data about the academic and career activities of 86 individuals from this group, of whom 77 (90%) attended community colleges in California, while the remaining 9 (10%) in Alabama, Arizona, Florida, Illinois, Massachusetts, Nevada, New York, and Washington. Almost half of CCI alumni attended six community colleges, listed here in descending order of attendance: Contra Costa College, Diablo Valley College, City College of San Francisco, College of Marin, College of San Mateo, and Hartnell College. As community college students taking lower -division courses, CCI interns are defined as taking coursework equivalent to that of first- or second-year undergraduates at baccalaureate granting institutions, even if they spent more than 2 years taking lower-division lower division coursework.

The second group, serving as the **comparison group**, consists of individuals who participated in the SULI program. To create as many similarities as possible between the study population and comparison group, we included *only* those SULI participants who were first- or second-year undergraduates at the time of their participation in SULI at LBNL. From the years 2009 to 2016, there were approximately 500 SULI participants at LBNL, and we had access to program records, which included information such as the institution in which they were enrolled at the time of their application to the SULI program, academic year of study, and previous higher education institutions attended. We were particularly interested in the last two categories, as they helped to determine which SULI alumni would be invited to participate in this study. Ultimately, the comparison group constructed for this study includes the 99 individuals who a) participated in the SULI program at LBNL from 2009 to 2016, b) were in the first or second year of their undergraduate studies at the time of their first internship term in SULI, and c) did not begin their studies at a community college. We collected data on the academic and career trajectories from 90 individuals who attended baccalaureate granting institutions across the U.S. The distribution of these institutions was as follows: 36 (40%) in California, 9 (10%) in Massachusetts, 7 (8%) in New York, 5 (6%) in Pennsylvania, 4 (4%) in North Carolina, and the remaining 29 (32%) in Colorado, Florida, Idaho, Illinois, Kansas, Kentucky, Michigan, Minnesota, Missouri, Nevada, New Jersey, Ohio, Oregon, Puerto Rico, Rhode Island, Tennessee, Texas, Vermont, Virginia, and Washington. Nearly 40% of SULI alumni attended the following seven universities, listed in descending order: University of California, Berkeley; University of Southern California; Princeton University; State University of New York at Stony Brook; University of California, Los Angeles; University of North Carolina, and University of Pennsylvania.

Of the CCI alumni who attended California Community Colleges, 60/77 (78%) attended a community college in the Bay Area, which consists of Alameda, Contra Costa, Marin, Napa, San Francisco, San Mateo, Santa Clara, Solano, and Sonoma counties (U.S. Census Bureau, 2011). At the time of data collection for this study, the LBNL main site was located in Alameda County, and off-site LBNL offices were located in Alameda and Contra Costa counties. Considering previous findings about community college student preference for academic/career opportunities within a “comfortable distance,” we compared the race and ethnicity of CCI participants with Bay Area census data from a similar time frame (Backes & Velez, 2015; Holland Zahner, 2022; Jabbar et al., 2017; Reyes et al., 2019), as shown in **Figure 1**. Additionally, we included SULI alumni data to compare how the CCI and SULI programs differ based on the race and ethnicity of participants.

**Figure 1.**
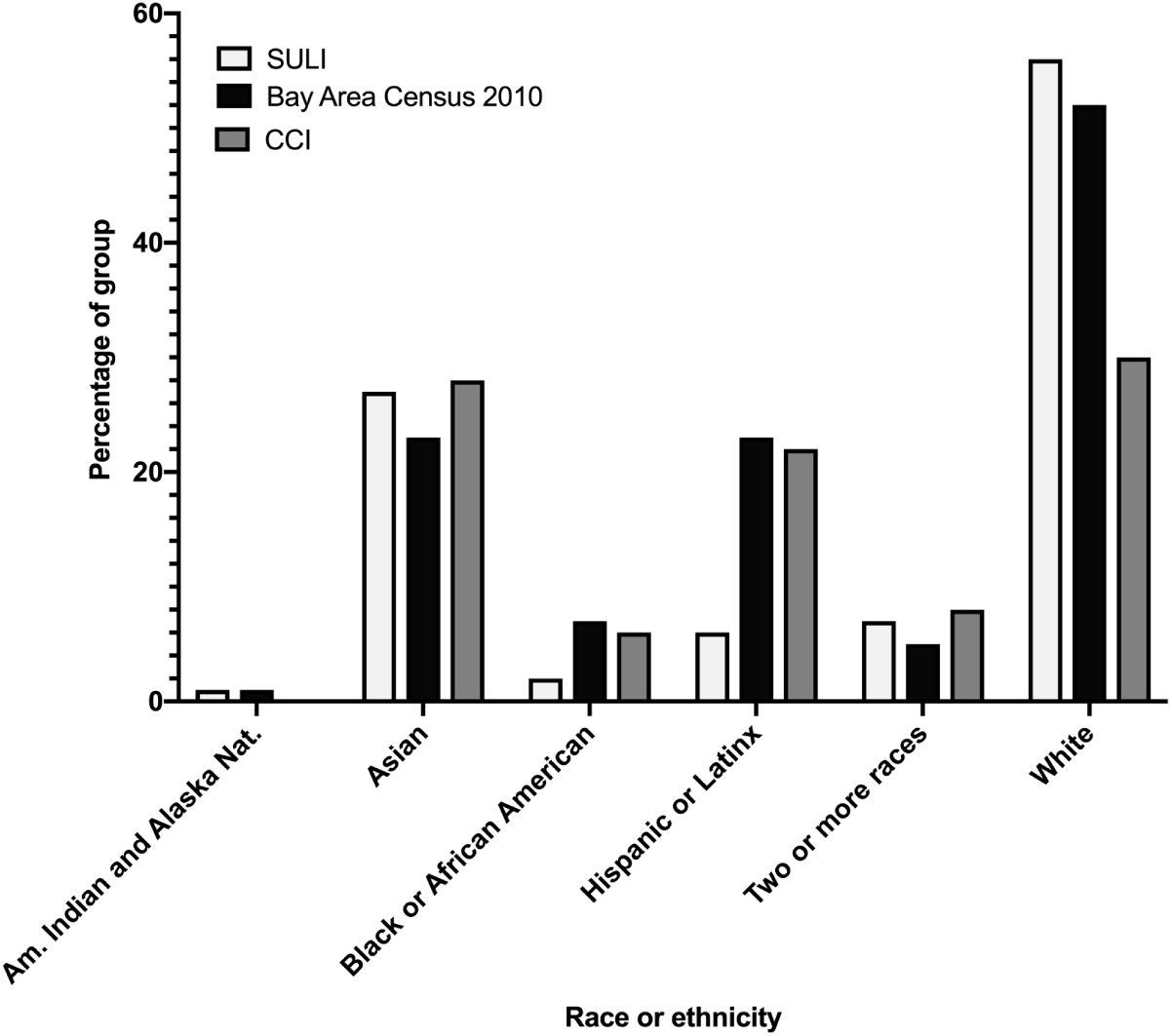
Comparison of Race/Ethnicity Distributions Between Study Participants and the 2010 Bay Area Census. *Note.* Based on the 2010 Census, the Bay Area population was 0.7% American Indian and Alaska Native, 23.3% Asian, 6.4% Black or African American, 23.5% Hispanic or Latinx, 0.6% Native Hawaiian and Other Pacific Islander, 10.8% some other race, 5.4% two or more races, and 55.6% White in 2010 (U.S. Census Bureau, 2011). During the 2009-2016 time frame, the CCI participants were 27.9% Asian, 5.8% Black or African American, 22.1% Hispanic or Latinx, 8.1% two or more races, 30.2% White, and 5.8% unknown; and the SULI participants were 1.1% American Indian and Alaska Native, 26.7% Asian, 2.2% Black or African American, 5.6% Hispanic or Latinx, 6.7% two or more races, and 52.5% White, and 2.2% unknown.

### Differences in school type and resources between groups

We constructed a comparison group (SULI alumni) that shares many similarities with the study population (CCI alumni), given the common features between the SULI and CCI programs. The primary difference is that SULI is open to all U.S. undergraduates and recent graduates, while CCI is specifically for U.S. community college students. Many alumni from both groups interacted at LBNL, attended the same group meetings, and/or worked in the same Mentor Groups.

However, there are some important differences between the two groups. For example, many U.S. colleges and universities, particularly those attended by SULI alumni, have large endowments that can be used to support many types of activities, including student financial aid and stipends, professional development programs, research supplies and facilities (ACE, 2021). Approximately 40% of public and 42% of private baccalaureate granting institutions have endowments larger than $50 million, and all of the SULI alumni in this study attended schools in this category (ACE, 2021; NACUBO, 2022). In contrast, only 6% of public and 5% of private baccalaureate granting institutions have endowments larger than $50 million, and 68 (76%) of SULI alumni attended schools in this category (ACE, 2021; NACUBO, 2022). Additionally, the eight baccalaureate granting institutions that make up the “Ivy League” in the U.S. – Brown University, Columbia University, Cornell University, Dartmouth College, Harvard University, Princeton University, University of Pennsylvania, and Yale University – are known for their high selectively and large endowments, and 14 (16%) of SULI alumni attended one of these prestigious and highly selective schools (U.S. News, 2022).

Moreover, of the SULI alumni who make up the comparison group, 75 (83%) attended schools in the 80th to 100th percentile of selectivity among all baccalaureate granting institutions, 73 (81%) attended schools classified as “research institutions,” 68 (76%) attended doctoral universities with “very high research activity,” 39 (43%) attended private schools, and 15 (17%) attended minority-serving institutions (ACE, 2022). In comparison, of the CCI alumni in this study, 74 (86%) attended minority-serving institutions, and none attended private schools, schools that would be considered to have high levels of “research activity,” or schools with large endowments (ACE, 2022). At the time of this writing, there is a permanent endowment of $25 million per year for the California Community College system, which is shared between 116 colleges across the state (Foundation for California Community Colleges, 2024).

### Data analysis

As described in the “Data collection” section, we collected academic and career activity data, which illustrates previous accomplishments, current employment, or current engagement in graduate studies. Information regarding the academic and career “trajectories” of both CCI and SULI alumni was organized by A.M., A.N.Z., L.E.C., J.J.S., and S.V.D. to show how many individuals from each group are on a specific career pathway: STEM, health, or non-STEM. This method of analysis was also used in our previous study on the CCI program (Coté et al., 2025). These categories were created to determine the proportion of each group that persisted in STEM, or is currently engaged in graduate-level training to support a STEM career. To assess whether there were statistically significant differences between outcomes (e.g., academic achievement, career activities, interest in working at LBNL in the future), for the CCI and SULI alumni group, we applied Fisher’s exact test to our data, and report two-tailed *p*-values (Armitage et al., 2013; Greenland, 1990). Fisher’s exact test was chosen because of the small sample size and the analysis of independent categorical data. This test is particularly appropriate for small sample sizes where the expected frequency of some categories is less than 5, as it provides an exact *p*-value rather than relying on approximations that may not be accurate with small sample sizes, Fisher’s exact test ensures more reliable results

Throughout this study, we refer to the “STEM workforce” and the majors, degrees, and pathways related to students preparing to enter this workforce. Although there are many studies published each year about STEM education, majors, careers, and workforce, these do not always specify which disciplines or professions are included in the “STEM” category (van den Hurk et al., 2019). To conduct our data analysis, we followed the guidance provided by the NASEM (2018) and National Science Board (2016), using what they refer to as “science and engineering occupations” (e.g., computer scientists, mathematicians, life scientists, physical scientists, engineers) and “science and engineering-related occupations” (e.g., laboratory managers, science and engineering teachers, lab technicians). A.M.B., A.N.Z., J.J.S., L.E.C., and S.V.D. identified and discussed the themes, and the data representation was refined for accuracy and clarity throughout the study.

To more simply illustrate the variety of academic and career trajectories of study participants, we present this data in Sankey diagrams, in **Figures 2 and 3**. The nodes — vertical bars — represent the number of individuals in each group, and represent the total number of individuals in the study and comparison groups, type of degree obtained, and current academic/career activity. The curved bands that run from the left-hand side of the diagram to the right-hand side are the flows, which are thicker when there are more individuals on that “path.” These flows show academic/career activities from undergraduate studies to entrance into the workforce or graduate studies.

**Figure 2.**
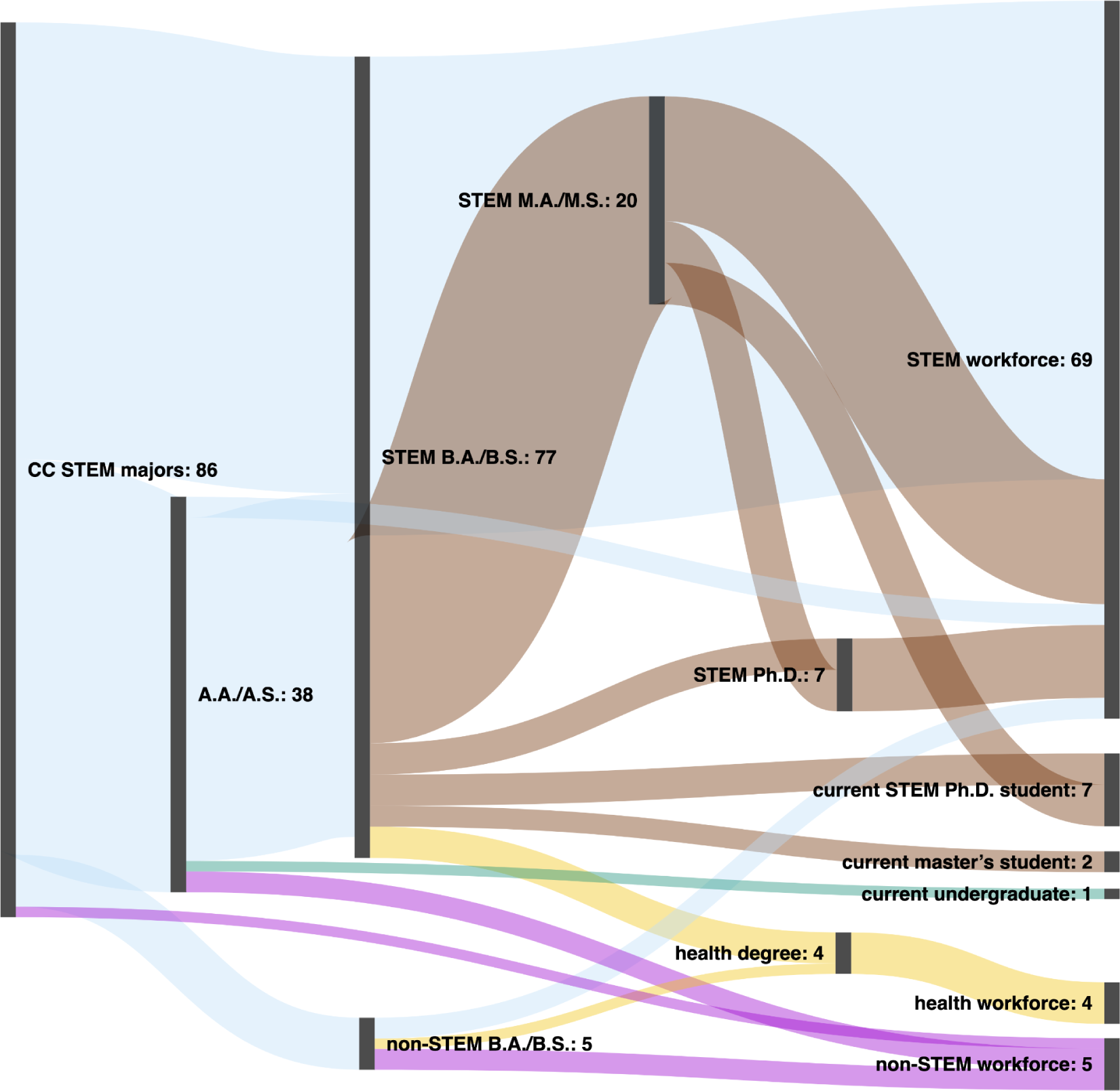
Academic and Career Trajectories of Community College Internship (CCI) Alumni. *Note.* In this Sankey diagram, the academic and career trajectories of Community College Internship (CCI) alumni (n=86) who participated in the program at LBNL between 2009 and 2016 are shown. The numbers next to each node indicate the number of individuals within the larger group of CCI alumni. For example, of the original 86 “STEM majors” (shown on the left side of the diagram), 38 obtained an A.A./A.S. degree, 77 obtained a STEM B.A./B.S. degree, and 5 obtained a non-STEM B.A./B.S. degree. This information is current through December 2021, which is 5 to 12 years after alumni participation in the CCI program.

**Figure 3.**
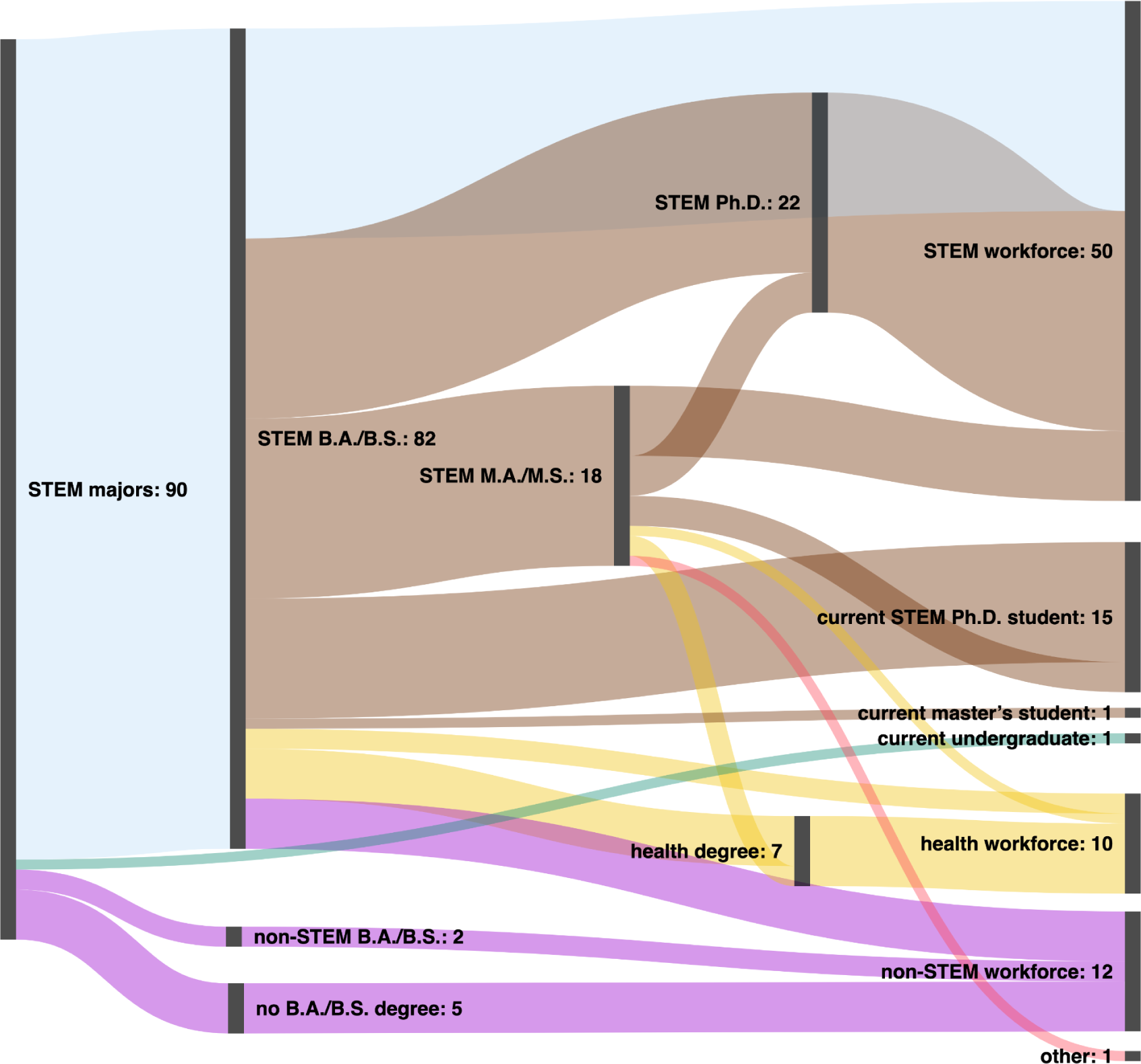
Academic and Career Trajectories of Science Undergraduate Laboratory Internship (SULI) Alumni. *Note.* In this Sankey diagram, the academic and career trajectories of Science Undergraduate Laboratory Internship (SULI) alumni (n=90) who participated in the program at LBNL between 2009 and 2016 are shown. The numbers next to each node indicate the number of individuals within the larger group of SULI alumni. For example, of the original 90 “STEM majors” (shown on the left side of the diagram), 82 obtained a STEM B.A./B.S. and 2 obtained a non-STEM B.A./B.S. This information is current through December 2021, which is 5 to 12 years after alumni participation in the SULI program.

## Limitations

### Selection bias

Selection bias is commonly described as a limitation of studies examining the impacts of internships and undergraduate research experiences (e.g., Provencher & Kassel, 2019; Taraban & Logue, 2012). It refers to a scenario where students who apply to or participate in a program are more likely to achieve academic and professional success than their peers. For example, students who have completed a STEM internship may, on average, have better grades, more relevant experience, and greater knowledge of STEM careers than their peers who did not participate in a STEM internship. We do not claim that participation in the CCI or SULI programs directly caused the academic or career activities of alumni after completing these programs. Instead, our study builds on an existing body of research linking participation in mentored internships and research experiences to increased confidence and interest in STEM, higher rates of STEM degree completion, and persistence in STEM fields (e.g., Eagan Jr et al., 2013; Gamage et al., 2021; Haeger et al., 2024; Hernandez et al., 2018; Higgins, 2013; Hirst et al., 2014; Jelks & Crain, 2020; Leggett-Robinson et al., 2015; Nerio et al., 2019; Prunuske et al., 2016).

### Demographic data collection

Although we have constructed a comparison group (SULI alumni) that is similar to the study population in many ways, there are some differences in terms of race/ethnicity. As shown in **Figure 1**, there were more Black/African American and Hispanic/Latinx participants in the CCI program, and more White participants in the SULI program during the 2009-2016 time frame. During the initial collection of demographic information from program participants, individuals were given the opportunity to select only one option related to race/ethnicity, including “two or more races,” which is a commonly used demographic category on many academic and professional records, surveys, etc. However, this category is not descriptive enough to allow for accurate separation of all participants into the commonly used “underrepresented minority” category, which generally includes individuals from the following groups: Alaska Native, Black/African American, Hispanic/Latinx, Native American, Native Hawaiian or other Pacific Islander (e.g., Robnett et al., 2015). Without this information, the individuals in the “two or more races” category are aggregated together, limiting the accuracy of data analysis related to race/ethnicity. One possibility is to include a checkbox question (for which respondents select one or more choices), allowing respondents to select the exact combination of categories they choose to report.

### Survey response rate

As part of the data collected for this study, we asked CCI and SULI alumni to respond to survey questions, but nonresponse error – where not all of the individuals in the study population have provided a response – is a potential limitation of this study (Ponto, 2015). Out of the individuals included in this study, 41 out of 86 (48%) CCI alumni and 57 out of 90 (63%) SULI alumni responded to survey questions about their interest level in working for LBNL or another DOE national laboratory or facility in the future. A 2019 study by Hendra and Hill suggests that a response rate of 80% (often used in federal reports) may not be necessary to ensure generalizable results. However, we recognize that response bias – individuals who choose to respond to a survey may be systematically different from those who do not respond – may limit the generalizability of our findings, because the smaller group of program alumni who responded to questions about their interest level in working for LBNL or another DOE national laboratory or facility may not be representative of the perspectives of all program alumni. Although not all of the program alumni responded to these survey questions, the groups that did respond had a demographic makeup similar to the full group (shown as “CCI DOE survey” and “SULI DOE survey” in **Table 1**). Our goal was to achieve the highest possible response rates, and the demographic similarities between groups reduce the likelihood that the responses would differ if every individual had responded (Hendra & Hill, 2019).

**Table 1.** Gender, Ethnicity, and Race of Study Participants.

| Demographics |  | CCI group |  | CCI DOE survey |  | SULI<sup>a</sup> group |  | SULI DOE survey |  |
| --- | --- | --- | --- | --- | --- | --- | --- | --- | --- |
|  |  | n | % | n | % | n | % | n | % |
| Total |  | 86 | 100 | 41 | 100 | 90 | 100 | 57 | 100 |
| Gender |  |  |  |  |  |  |  |  |  |
|  | Female | 27 | 31 | 18 | 44 | 36 | 40 | 21 | 37 |
|  | Male | 58 | 67 | 22 | 54 | 54 | 60 | 36 | 63 |
|  | Unknown<sup>b</sup> | 1 | 1 | 1 | 2 | 0 | 0 | 0 | 0 |
| Ethnicity/race |  |  |  |  |  |  |  |  |  |
|  | Asian | 24 | 28 | 12 | 29 | 24 | 27 | 9 | 16 |
|  | Black or African American | 5 | 6 | 0 | 0 | 2 | 2 | 1 | 2 |
|  | Hispanic or Latinx | 19 | 22 | 9 | 22 | 5 | 6 | 4 | 7 |
|  | Native American | 0 | 0 | 0 | 0 | 1 | 1 | 0 | 0 |
|  | Two or more races | 7 | 8 | 3 | 7 | 6 | 7 | 4 | 7 |
|  | White | 26 | 30 | 15 | 37 | 50 | 56 | 38 | 67 |
|  | Unknown<sup>b</sup> | 5 | 6 | 2 | 5 | 2 | 2 | 1 | 2 |

*Note.* This table includes demographic information for the Community College Internship (CCI) and Science Undergraduate Laboratory Internship (SULI) participants who completed their program at LBNL between 2009 and 2016. <sup>a</sup> This is a subset of all 2009-2016 SULI participants at LBNL. This group includes only those who were first- and second-year undergraduates during their first SULI term and did not attend a community college at any time prior to their participation in SULI. <sup>b</sup> For both gender and ethnicity/race data, the “unknown” group includes both “decline to state” responses and missing data.

## Results

### Academic and career trajectories for CCI alumni

We collected data about the academic and career activities of 86 CCI alumni, between 5 and 12 years after their participation in the CCI program at LBNL (**Table 2**). As shown in **Figure 2**, all (100%) of these 86 CCI alumni were “STEM majors” attending a community college at the time of their participation, 38 (44%) obtained one or more A.A./A.S. degrees, 77 (90%) obtained a STEM B.A./B.S., 5 (6%) obtained a non-STEM B.A./B.S. degree, and 1 (1%) is currently an undergraduate student. Nearly all CCI alumni (81; 94%) transferred from a community college to a baccalaureate granting institution; 46 (53%) transferred *without* first obtaining an A.A./A.S. degree; and 35 (41%) transferred *after* obtaining one or more A.A./A.S. degrees. After completing their CCI internship term at LBNL, 14 (16%) alumni participated in the SULI program at LBNL, 5 (6%) participated in a different internship at LBNL, and 9 (10%) participated in an internship at a different DOE national laboratory.

**Table 2.** Current Academic and Career Activities of Study Participants.

| Academic or<br>career activity | CCI<br>alumni |  | SULI<br>alumni |  | CCI/SULI alumni<br>combined |  |
| --- | --- | --- | --- | --- | --- | --- |
|  | *n* | % | *n* | % | *n* | % |
| Total | 86 | 100 | 90 | 100 | 176 | 100 |
| Undergraduate or post-baccalaureate<br>program (STEM) | 1 | 1 | 1 | 1 | 2 | 1 |
| Master's program (STEM) | 1 | 1 | 0 | 0 | 1 | 0.6 |
| Master's program (other) | 1 | 1 | 1 | 1 | 2 | 1 |
| Ph.D. program (STEM) | 7 | 8 | 14 | 16 | 22 | 12.5 |
| Ph.D. program (other) | 0 | 0 | 1 | 1 | 1 | 0.6 |
| STEM workforce*** | 69 | 80 | 50 | 56 | 118 | 67 |
| Health workforce | 4 | 5 | 10 | 11 | 14 | 8 |
| Non-STEM workforce | 5 | 6 | 12 | 13 | 17 | 10 |
| Other | 0 | 0 | 1 | 1 | 1 | 0.6 |

*Note.* The academic or career activities of participating individuals may be described by more than one category. For example, two CCI alumni joined the workforce and started an academic program without leaving their professional positions. Fisher's exact test was applied to determine if there was a significant difference between the number of alumni from each program (first and second columns in Table 2) associated with each academic/career activity. Two-tailed P-values are labeled as follows: \*, $p < 0.1$ significance level; \*\*, $p < 0.05$ significance level; \*\*\*, $p < 0.01$ significance level. This information is current through December 2021, which is 5 to 12 years after alumni participation in the CCI and SULI programs.

Of the 77 CCI alumni who obtained a STEM B.A./B.S., 73 (95%) entered a STEM field through the workforce or graduate studies, 4 (5%) joined the health workforce and none (0%) joined the non-STEM workforce. In comparison, of the 5 individuals who obtained a non-STEM B.A./B.S. degree, 2 (40%) joined the non-STEM workforce, 2 (40%) joined the STEM workforce, and 1 (20%) joined the health workforce. Of this group of CCI alumni who obtained a STEM B.A./B.S., 20 (26%) obtained a STEM M.A./M.S. degree, 2 (3%) are currently enrolled in a STEM M.A./M.S. degree program, 7 (9%) obtained a STEM Ph.D. degree, 7 (9%) are currently enrolled in a STEM Ph.D. program, 3 (4%) obtained a health degree (e.g., D.D.S., M.D., Pharm.D.), 65 (84%) are currently part of the STEM workforce, and 3 (4%) are currently part of the health workforce. Finally, 5 (4%) CCI alumni graduated with a B.A./B.S. degree in a non-STEM subject *and* joined the non-STEM workforce before enrolling in a community college to take STEM coursework.

### Academic and career trajectories for SULI alumni

We collected data about the academic and career activities of 90 SULI alumni (who were first- or second-year undergraduates at the time of their internship) between 5 and 12 years after they participated in the SULI program at LBNL. As shown in **Figure 3**, of the 90 SULI alumni in this study, 82 (91%) obtained a STEM B.A./B.S., 2 (2%) obtained a non-STEM B.A./B.S. degree, 5 (6%) did not obtain a B.A./B.S. degree in any subject, and 1 (1%) is currently an undergraduate student. Of the 82 SULI alumni who obtained a STEM B.A./B.S. degree, 66 (80%) entered a STEM field through the workforce or graduate studies, 10 (12%) joined the health workforce, and 5 (6%) joined the non-STEM workforce. All (100%) of the individuals who obtained non-STEM B.A./B.S. degrees *or* did not obtain B.A./B.S. degrees have entered the non-STEM workforce. Of the 82 SULI alumni who obtained a STEM B.A./B.S., 18 (22%) obtained a STEM M.A./M.S. degree, 1 (1%) is currently enrolled in a STEM M.A./M.S. degree program, 22 (27%) obtained a STEM Ph.D. degree, 15 (18%) are currently enrolled in a STEM Ph.D. program, 7 (9%) obtained a health degree (e.g., D.D.S., M.D., Pharm.D.), 50 (61%) are currently part of the STEM workforce, 10 (12%) are currently part of the health workforce, and 5 (6%) are currently part of the non-STEM workforce.

### Comparing the trajectories of CCI and SULI alumni

As described in the “Data analysis” section, we used three categories to identify those individuals who are currently working and/or attending school in a discipline related to STEM, health, or non-STEM. As opposed to the current academic and career activities data shown in **Table 2**, the data in **Figure 4** combine the data from individuals in the workforce with those enrolled in graduate programs to produce STEM, health, or non-STEM “career pathways” categories. For example, the number of individuals working in the health workforce as medical staff, dentists, or physicians have been combined with the number of individuals currently enrolled in nursing, dental, or medical degree programs to produce a single “health career pathway” group. Based on the “career pathways” data (see **Table S3** for details), a statistically significant greater proportion of **CCI alumni (88%) than SULI alumni (71%) are currently on a STEM career pathway** (*p*-value = 0.0051; odds ratio = 3.088 with 95% CI of 1.439 to 6.682). In other words, CCI alumni are 3 times more likely to pursue a STEM career pathway than SULI alumni. We found that 6% of CCI alumni and 13% of SULI alumni are currently on a health career pathway, and 6% of CCI alumni and 16% of SULI alumni are currently on a non-STEM career pathway (*p*-value = 0.0509; odds ratio = 0.3351 with 95% CI of 0.1286 to 0.9344).

**Figure 4.**
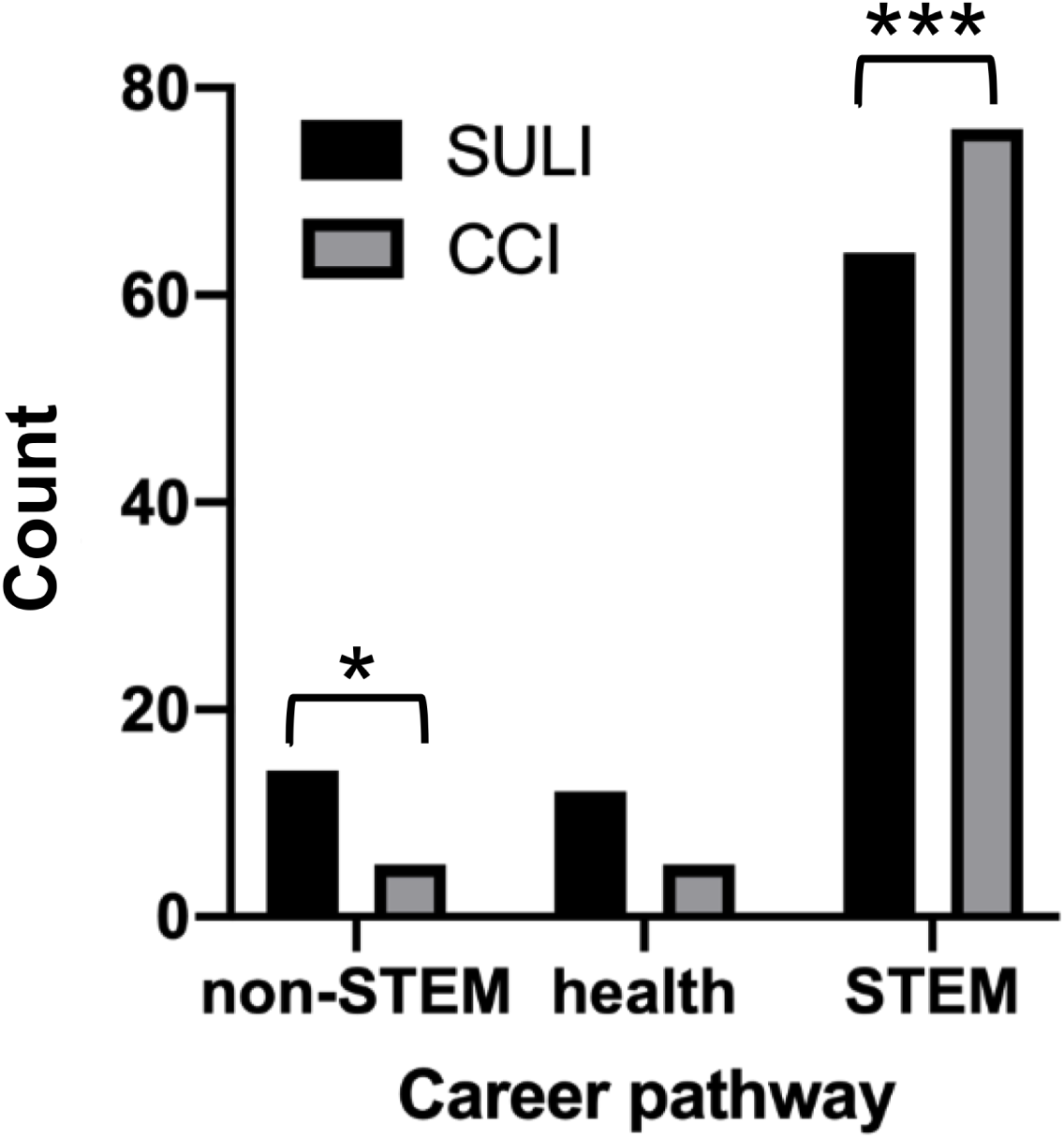
Career Pathways for Study Participants by Program. *Note.* For SULI alumni – who were first- or second-year undergraduates attending baccalaureate granting institutions at the time of their internship – 64 (71%) are on a STEM career pathway, 12 (13%) are on a health career pathway, and 14 (16%) are on a non-STEM career pathway. For CCI alumni – who were community college students during the internship – 76 (88%) are on a STEM career pathway, five (6%) are on a health career pathway, and five (6%) are on a non-STEM career pathway. Fisher’s exact test was applied to determine if there was a significant difference between the number of alumni from each program associated with each career pathway. Two-tailed P-values are labeled as follows: *, *p* < 0.1 significance level; **, *p* < 0.05 significance level; ***, *p* < 0.01 significance level. This information is current through December 2021, which is 5 to 12 years after alumni participation in the CCI and SULI programs.

Considering those individuals who are currently employed, we found that a statistically significant greater proportion of **CCI alumni (80%) than SULI alumni (56%) have joined the STEM workforce** (*p*-value = 0.0007; odds ratio = 3.247 with 95% CI of 1.660 to 6.148). In other words, CCI alumni are 3.2 times more likely to join the STEM workforce than SULI alumni. Additionally, 5% of CCI alumni and 11% of SULI alumni have joined the health workforce, and 6% of CCI alumni and 13% of SULI alumni have joined the non-STEM workforce.

As shown in **Table S3**, we collected data about the academic achievements of program alumni and found that 90% of CCI alumni and 91% of SULI alumni have completed a STEM B.A./B.S. degree, and 23% of CCI alumni and 20% of SULI alumni have completed a STEM master’s degree. We found that a statistically significant lesser proportion of CCI alumni (8%) than SULI alumni (24%) have completed a STEM Ph.D. degree (*p*-value = 0.0042; odds ratio = 0.2739 with 95% CI of 0.1071 to 0.6949). In other words, SULI alumni are 3.7 times more likely to obtain a STEM Ph.D. than CCI alumni. Of those who completed degrees outside of STEM: 6% of CCI alumni and 2% of SULI alumni have completed a non-STEM B.A./B.S. degree; and 5% of CCI alumni and 7% of SULI alumni have completed a health or non-STEM graduate degree.

### Alumni experience and interest in working at national laboratories

The CCI and SULI programs are part of a suite of programs hosted at national laboratories designed to expose students and recent graduates to career opportunities at these institutions (DOE, 2022). Alumni from these programs are thus considered to be candidate pools from which national laboratories can “recruit new permanent hires” (DOE SC, 2020). As described in the “Limitations” section, a smaller set of program alumni responded to survey questions about their interest in working at LBNL or another DOE national laboratory or facility in the future. Since their participation in the internship at LBNL, 19/41 (46%) of CCI alumni and 17/57 (30%) of SULI alumni worked at LBNL in some capacity. A statistically significant greater proportion of CCI alumni (26/41, 63%) than of SULI alumni (21/57, 37%) worked at *any* of the DOE national laboratories or facilities in some capacity (*p*-value = 0.0137; odds ratio = 2.971 with 95% CI of 1.250 to 7.041). In other words, CCI alumni were 3 times more likely than SULI alumni to work at a DOE national laboratory or facility in the years following their internship. Most commonly, alumni worked at a DOE national laboratory (including LBNL) or facility after completing an additional internship program.

Currently, 8/41 (20%) of CCI alumni and 5/57 (9%) of SULI alumni who responded to these survey questions work at one of six different national laboratories. Their roles include analysts, engineers, postdoctoral scholars, research assistants, research associates, and scientists. They work in various disciplines, including chemical engineering, chemistry, environmental science, materials science, mechanical engineering, nuclear engineering, nuclear science, operations, and physics. Additionally, we learned that a statistically significant greater proportion of of CCI alumni (33/41, 80%) than of SULI alumni (29/57, 51%) are interested in working at LBNL in the future (*p*-value = 0.0031; odds ratio = 3.983 with 95% CI of 1.596 to 10.01). In other words, CCI alumni were 4 times more likely to want to work at LBNL in the future than SULI alumni. Both CCI alumni (15/41, 37%) and SULI alumni (24/57, 42%) were interested in working at a *different* DOE national laboratory in the future. Combining the CCI and SULI groups together, **13/98 (13%) of alumni currently work at one of the national laboratories**, and 64/98 (65%) of alumni are *interested* in working within the DOE complex now or in the future.

## Discussion

### Outcomes reveal CCI to be a “high impact” program

Many individuals who enroll in community colleges are hoping to gain new knowledge, develop valuable skills, and advance their career, which are aligned with our previous findings about outcomes for alumni of the CCI program at LBNL (Coté et al., 2025; Torpey-Saboe & Clayton, 2023). Thus, there is great potential for programs like CCI to *improve* academic and career outcomes for participants from all backgrounds. At the time of their participation in CCI at LBNL, the study population (CCI alumni) attended public community colleges with little to no opportunity for undergraduates to conduct research on campus. Many individuals in the comparison group (SULI alumni) were students at some of the most selective, well-funded, and research intensive baccalaureate granting institutions. Based on the differences between the schools attended by members of each group – especially in terms of resources available to support student professional development and success – we would expect to see major differences in academic/career activities between groups. However, our study found high rates of degree attainment, retention in STEM majors, and entry into the STEM workforce from both CCI and SULI alumni. We acknowledge that there are likely many differences between the two groups of students we studied, but believe that the results from our previous study about the impacts of CCI at LBNL (Coté et al., 2025) coupled with the career pathway data presented in the current study strongly illustrate the benefits of offering discipline-specific professional development opportunities to community college students.

### CCI alumni graduated at rates that were higher than expected for community college STEM majors

Multiple studies have shown that students who enroll in community colleges to obtain a STEM degree often do *not* have the support to accomplish this goal, especially for Alaska Native, Black/African American, Hispanic/Latinx, Native American, female, first-generation to college, and low-income students (e.g., CCSSE, 2021; Chen, 2013; Hill et al., 2010; Huang et al., 2000; Kokkelenberg & Sinha, 2010; Sansing-Helton et al., 2021; Torpey-Saboe & Clayton, 2023; Varty, 2022). More than two-thirds of STEM majors attending community colleges end up “leaving STEM” by selecting a non-STEM course of study or dropping out of their undergraduate institution (Chen, 2013). Based on what is known about community college student trajectories, we found that CCI alumni transferred and graduated with bachelor’s degrees at *higher* rates than expected. Data collected about students who attended U.S. community colleges during the time frame we studied revealed that 20-30% graduated with an associate degree and 9-16% graduated with a bachelor’s degree within 6 years (CCSSE, 2021; Horn & Skomsvold, 2011; Juszkiewicz, 2020; Sansing-Helton et al., 2021; De Brey et al., 2021). In comparison, 44% of the CCI alumni in our study graduated with an associate degree, and 96% graduated with a bachelor’s degree. In our previous study about the CCI program, we learned that one-third of alumni were *more* interested in graduating with a bachelor’s degree after their participation in the CCI program at LBNL (Coté et al., 2025). Aligned with previous studies, participation in STEM technical/research experiences can provide academic and attitudinal benefits to community college participants (e.g., Gamage et al., 2021; McIntyre et al., 2020; Nerio et al., 2019).

### “Career changers” leverage community colleges and the CCI program

Approximately 62% of people who graduate from high school begin their undergraduate studies at a college or university in the same calendar year (NCES, 2024). However, there are many pathways into higher education, and some individuals choose to attend college later in life. The average age of enrolled students at U.S. community colleges is 27 years, and some students possess a non-STEM bachelor’s degree before enrolling in community college with the goal of entering the STEM workforce (AACC, 2024; Allison et al., 2022). Similarly, our study found that 5 (4%) CCI alumni graduated with a non-STEM B.A./B.S. degree *and* joined the non-STEM workforce before enrolling in a community college to take STEM coursework. Referred to as “career changers” in the higher education literature, individuals who enter their undergraduate studies at community colleges with previous professional experiences could prove to be *valuable* assets to the U.S. STEM workforce. Further long-term exploration of the academic and career activities of program alumni will determine if this is the case at other institutions that host CCI or similar programs for community college students taking coursework in STEM disciplines.

### Most CCI alumni are on STEM career pathways and interested in working in the DOE complex

In the U.S., the majority of STEM professional development opportunities for undergraduate students are a) hosted by baccalaureate granting institutions, or b) involve partnerships between community colleges and baccalaureate granting institutions located near each other (Draganov et al., 2023; Nerio et al., 2019). Our findings show that – as compared to students attending baccalaureate granting institutions – community college students who engage in STEM professional development activities are likely to persist in STEM careers at similar rates. This finding aligns with previous studies that have recommended partnerships between community colleges and industry partners to support “gainful employment for students” (e.g., Atwell et al., 2022; D’Amico et al., 2019).

A number of reports have been produced to document discipline- or industry-specific workforce needs in the U.S., analyze the gaps in workforce development, and identify strategies to address these (e.g., McDowell et al., 2024; Paz-Soldan et al., 2025; The Fusion Cluster, 2023). Many of these highlight the relatively unknown relationship between internships and other training programs and alumni retention in STEM or industries. Our data about alumni interest in working at a DOE national laboratory are relevant to efforts across the DOE complex to train students and recent graduates through internships and retain some of these alumni as permanent staff at national laboratories and facilities (DOE, 2022; DOE SC, 2020). Based on survey data collected about the career activities and perspectives of past program participants, 65% of alumni expressed interest in working at a DOE national laboratory or other DOE facility, but only 13% of alumni are currently employed at one of these institutions. Notably, after the completion of the internship, CCI alumni were three times *more* likely than SULI alumni to have worked at a DOE national laboratory or facility. Additionally, CCI alumni were four times *more* likely than SULI alumni to be interested in working at LBNL in the future. We recommend this type of analysis be repeated in the future, to determine if similar patterns are observed with a larger group and/or with alumni of discipline- or industry-specific programs. Although we cannot foresee future career activities of the CCI and SULI alumni in this study, our findings appear to conflict with a previous estimate that “roughly 50 percent of program participants eventually work at a national laboratory” (Foltz et al., 2011). Thus, there is an opportunity to engage with this population about jobs at national laboratories through long-term communication and engagement to *increase* the number of alumni working in the DOE national laboratory system. That said, we find that CCI at LBNL has achieved the program goal of supporting participants to enter careers in STEM.

### Most SULI alumni are on STEM career pathways and are more likely to obtain a Ph.D. than CCI alumni

A wide variety of studies about students attending baccalaureate granting institutions have found that gains related to retention in STEM (e.g., graduation rates, entry into the STEM workforce, graduate school attendance) are supported through participation in research experiences, especially for students from groups historically excluded from STEM fields (Carpi et al., 2017; Estrada et al., 2018; Rodenbusch et al., 2016; Sadler et al., 2010). Our findings support this, and we found that most SULI alumni had either entered the STEM workforce and/or were actively completing their graduate studies in a STEM discipline. Further, the SULI learning environment – DOE national laboratories – and who choose to serve as mentors may have an impact on participants’ graduate school participation in the years following the program. Previous studies have shown that, when compared to students with similar background and achievement levels, undergraduates who participate in research experiences are more likely to complete a bachelor’s degree and pursue a Ph.D., especially when mentors discuss graduate school and career options (Camacho et al., 2021; Haeger et al., 2024; Romero et al., 2023; Trott et al., 2020; Wilson et al., 2018).

As described in the Introduction section, a previous study examined the programmatic elements of SULI, and reported some findings related to alumni perspectives. They administered surveys to 316 individuals who completed the SULI program between 2001 and 2007 at a variety of DOE national laboratories, and received responses from 70 individuals (Foltz et al., 2011). As a result, they found that 66% of SULI alumni were interested in obtaining “a doctoral degree or other terminal degree in a STEM discipline.” In comparison, our findings about SULI alumni indicated that 37 (41%) either completed or were currently pursuing a Ph.D. degree in STEM. Although the language used in the study by Foltz and colleagues may have included other terminal degrees (e.g., M.D., D.O., or D.D.S.), the results of these two studies are generally complementary regarding the interest of SULI alumni to pursue a doctoral degree in a STEM discipline. Additionally, though many alumni from both programs pursued graduate studies, we found that SULI alumni were nearly four times *more* likely than CCI alumni to complete a Ph.D. in a STEM discipline. In summary, SULI at LBNL has achieved the program goal of supporting participants to pursue STEM careers.

Scholars have identified several factors that may impact the representation of community college alumni in doctoral programs, often referred to as the “community college to Ph.D.” pipeline (e.g., Nguyen et al., 2024). These include academic advising that fails to introduce graduate school as an option, the lack of technical/research experiences available to community college students, challenges after transferring to baccalaureate granting institutions, and biases against community college students embedded in graduate school admissions processes (Kogan et al., 2015; Hewlett, 2016; Nguyen et al., 2024). Addressing these issues is beyond the scope of this study, but we believe they are worthy of future investigation.

## Recommendations for future studies

### Understand the role of “graduation” as a concept for community colleges

Many studies and national reports have used retention and graduation rates to determine the success of a community college or baccalaureate granting institution and/or to award funding or other forms of support (Cohen & Kelly, 2019). Although these metrics can be useful in understanding student success at an undergraduate institution, many researchers neglect to distinguish between community college students who "drop out" of school (i.e., leave their undergraduate studies without obtaining any degree), transfer between community colleges, and those who transfer without first obtaining an associate degree (Bahr, 2012; Porter, 2003). Additionally, it is common for students at community colleges to switch between “full-time” and “part-time” enrollment, which can have large impacts on the rates of graduation within a 3-year period of time (Cohen & Kelly, 2019). In our study, if we did not collect data about transferring (versus graduation), and/or the students in our study population were not tracked *beyond* their attendance in community college, it would be possible to conclude that 46 (53%) of CCI alumni “did not graduate.” Instead, we know that 81 (94%) of CCI alumni completed their studies at a community college (with or without obtaining an associate degree), and 78 (96%) of these transfer students completed their studies at a baccalaureate granting institution (by obtaining one or more bachelor’s degrees). We recommend considering the nuances inherent in studying the concepts of retention and graduation rates at different institution types.

### Study academic/career outcomes of community college students to align with workforce goals

As described in the previous section, we found that nearly all community college students who completed the CCI program at LBNL transferred to a baccalaureate granting institution and obtained a bachelor’s degree. If institutions would like to encourage more community college students to enter the middle-skill STEM workforce, this may require an examination of the Mentor Groups participating in the program, to ensure that professionals with relevant technical expertise are working with student participants. For example, many jobs in the energy workforce do not require a bachelor’s degree, and U.S. community colleges are likely to play an important role in providing education and training to individuals who would like to enter this sector (EWAB, 2024; NIST, 2023). Strategic recruitment of Mentor Groups with projects that will provide students with training in skills aligned with entry into the middle-skill STEM workforce, such as computer-aided design (CAD), cybersecurity, digital imaging, information technology, manufacturing, mechanical systems, microscopy equipment maintenance, operations management, procurement, robotics, vacuum systems, vehicle technology, etc. can help to achieve this goal (Moore et al., 2021; NASEM, 2017a; NLDC, 2024; The Fusion Cluster, 2023). However, if institutions intend to support community college students in obtaining bachelor’s degrees in preparation for entry into the STEM workforce, the current model used at LBNL is likely to be effective in achieving these goals. In summary, we recommend that host institutions and funding agencies consider the desired academic and career activities of its program alumni. Data collected from program alumni about these activities can be used by institutions to determine how to modify programs like CCI to better align with their goals.

### Maintain contact with program alumni

Considering the dearth of research on the topic, we believe that this study is a first step in understanding long-term outcomes for community college students, and how these compare to students from baccalaureate granting institutions. These findings are helpful in understanding the academic and career activities of alumni years after their participation in CCI or SULI, but – like many studies – this is not a “complete picture” of their activities. We were unable to communicate with every individual who completed these programs at LBNL during the 2009-2016 time frame, because much of the contact information we had on file was outdated at the time of our data collection efforts. This was especially true for those individuals who participated in the program in earlier years (2009-2011). Some individuals were only accessible to us through *temporary* email addresses associated with a college or university that expired once those students were no longer enrolled. Inspired by the National Science Foundation Survey of Earned Doctorates (an annual exit survey of all U.S. doctoral graduates), we recommend that programs administer post-surveys to past participants to a) collect personal email addresses, b) track the completion of higher education degrees, and c) gather information about academic and career activities. To further enhance access to information about academic and career trajectories, we encourage programs to learn more about how their study populations use professional networking websites, such as LinkedIn.

### Provide opportunities for community college students in STEM

Many individuals who complete some portion of their undergraduate studies at a community college join the STEM workforce after graduation, in middle-skill jobs, or those that require a higher degree (Belfield & Brock, 2023). Unfortunately, only a small percentage of the credentials offered by community colleges are “in demand by industry,” which has contributed to a shortage of skilled workers in the STEM workforce (NASEM, 2023). The implementation of research/technical experiences for (e.g., faculty-led research projects, course-based research experiences, internships) are ways to increase the number of individuals with the necessary skills and knowledge needed to enter the STEM workforce after their undergraduate studies. However, in the U.S., there are far fewer of these opportunities available to students at community colleges than those at baccalaureate granting institutions. For example, in 2022 there were approximately 100 CCI interns and 1,000 SULI interns working at one of the participating DOE national laboratories and facilities across the country.

Across the DOE national laboratory complex, there are other workforce development strategies that engage community colleges. For example, Los Alamos National Laboratory (LANL) partnered with Santa Fe Community College to produce curriculum and activities related to metalworking in alignment with LANL staffing needs (Santa Fe Community College, 2020). Similarly, Oak Ridge National Laboratory (ORNL) partnered with Roane State Community College on initiatives to provide coursework and skill development in nuclear technology, chemical engineering, and project management aligned with ORNL workforce needs (Butler & McKinney, 2023; Shoemaker, 2024). The STEM Core Community College program at SLAC National Accelerator Laboratory has provided community college students with research/technical internships (SLAC National Accelerator Laboratory, 2025). In the future, we hope to see community colleges, DOE national laboratories, and industry partners working together to provide additional STEM-based professional development opportunities for community college students.

## Conclusions

Our primary goal for this study was to examine the academic and career trajectories of CCI and SULI alumni, and determine the success of these programs to support entry into STEM careers. We found that most alumni from *both* the CCI and SULI programs were retained in STEM through degree programs and/or the workforce. Our work supports previous scholarship that undergraduates who participate in mentored technical/research experiences are more likely to *stay* in a STEM major, graduate with one or more STEM degrees, and enter a career in STEM. Collectively, 80% of SULI/CCI alumni are on a “STEM career pathway,” 77% completed a STEM bachelor’s degree, 20% completed a STEM master’s degree, and 16% completed a STEM Ph.D. Overall the current study aligns with previous findings that undergraduates who complete STEM technical or research experiences are more likely to *stay* in a STEM major, transfer to a baccalaureate granting institute (if attending a community college), graduate with one or more STEM degrees, and enter a career in STEM (e.g., McDaniel & Van Jura, 2022).

Many federal calls to action, DOE reports and strategic plans, and STEM education studies about professional development opportunities for undergraduates include recommendations to collect data about outcomes related to retention in STEM majors, graduation rates, and entry into the STEM workforce. However, most studies do not distinguish between outcomes for students who began their studies at a community college versus a baccalaureate granting institution. Beyond this, what is publicly known (i.e., published in scholarly journals) about the academic and career trajectories of STEM internships and research experiences almost always involves programs hosted at baccalaureate granting institutes. Coupled with our previous study (Coté et al., 2025), we believe that our findings about the career pathways of alumni make a strong case for the benefits of offering discipline-specific professional development opportunities to community college students. We found that U.S. community college students who complete professional development activities in STEM (such as the CCI program) are likely to complete their academic degrees and pursue STEM careers at rates comparable to those of students attending well-resourced baccalaureate granting institutes. Thus, we encourage educational researchers and scientists from other disciplines to partner with each other to conduct and *publish* work about internships and research experiences hosted at community colleges, national laboratories, companies, and other types of institutions (Collins, 2023; Hora et al., 2017; Lucero et al., 2021). We also encourage partnerships between community colleges and other institutions to provide community college students with opportunities to learn about careers in STEM.

## Acknowledgments

We are grateful to Laura Armstrong, Katie Blackford, Dianna Bolt, Sara Bourne, Beatriz Brando, Sean Burns, Chris Byrne, Christel Cantlin, Rachel Carl, Joseph Crippen, Michelle Douskey, Faith Dukes, Colette Flood, Ping Ge, Lloyd Goldwasser, Kris Gutiérrez, Ibrahim Hajar, Nakeiah Harrell, Max Helix, Abigail Hinojosa, Lady Idos, Bill Johansen, Terry Johnson, Geri Kerstiens, Kelsey Miller, Rey Morales, Colette Patt, Laura Pryor, Michael Ranney, Alexis Shusterman, Renée Schwartz, Christina Teller, Matt Traxler, Yukiko Watanabe, Michelle Wilkerson, Brieanna Wright, Marissa Yáñez, and the Baranger (ChemEd) Research Group at the University of California, Berkeley for their assistance, support, and feedback on this work.

## Disclosure Statement

The authors report there are no competing interests to declare.

## Funding

This material is based upon work supported by the National Science Foundation Graduate Research Fellowship Program under Grant No. DGE 1106400; Wheelhouse: The Center for Community College Leadership and Research at the University of California, Davis; the Berkeley School of Education Barbara Y. White Fund; and Workforce Development & Education at LBNL.

## Data Availability

The data set collected for this study contains detailed information about academic and career activities that could compromise the privacy of research participants. So, the full dataset generated for the current study is not publicly available. However, a selection of data that support the findings of this paper will be available from the corresponding author upon reasonable request.

## Supplemental materials

**Table S1.**
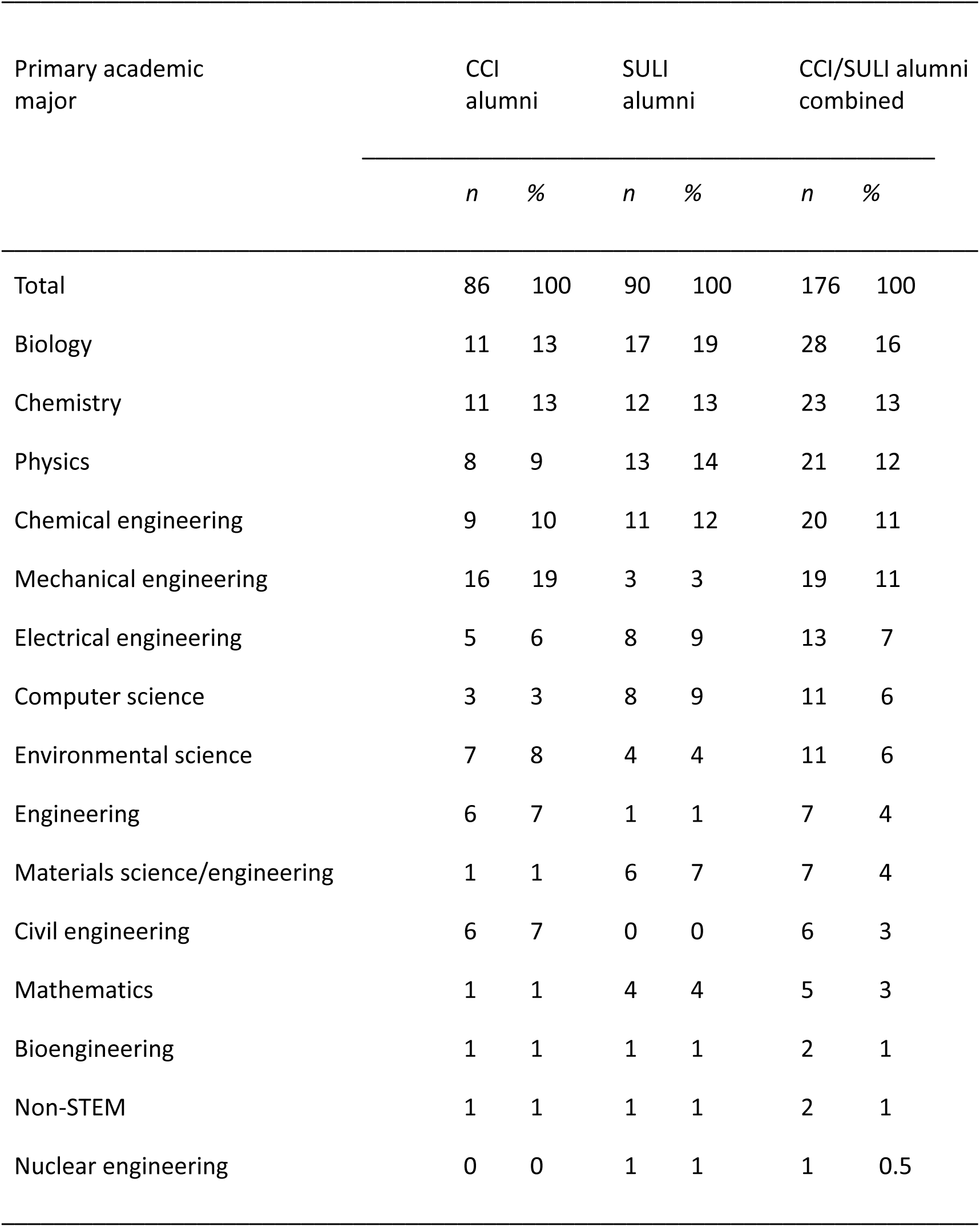
Primary Academic Major of Study Participants.

**Table S2.** Current Jobs held by Study Participants.

| Current job | CCI alumni |  | SULI alumni |  | CCI/SULI alumni combined |  |
| --- | --- | --- | --- | --- | --- | --- |
|  | *n* | % | *n* | % | *n* | % |
| Total | 86 | 100 | 90 | 100 | 176 | 100 |
| Research or laboratory assistant <sup>a</sup> | 8 | 9 | 16 | 18 | 24 | 14 |
| Chemical engineer | 9 | 10 | 10 | 11 | 19 | 11 |
| Civil or mechanical engineer | 17 | 20 | 1 | 1 | 18 | 10 |
| Software engineer | 6 | 7 | 8 | 9 | 14 | 8 |
| Physicist | 3 | 3 | 8 | 9 | 11 | 6 |
| Chemist | 3 | 3 | 6 | 7 | 9 | 5 |
| Postdoctoral scholar | 1 | 1 | 8 | 9 | 9 | 5 |
| Engineer | 6 | 7 | 1 | 1 | 7 | 4 |
| Biologist | 3 | 3 | 3 | 3 | 6 | 3 |
| Clinical scientist | 3 | 3 | 3 | 3 | 6 | 3 |
| Electrical engineer | 0 | 0 | 5 | 6 | 5 | 3 |
| Environmental scientist | 5 | 6 | 0 | 0 | 5 | 3 |
| Physician | 2 | 2 | 3 | 3 | 5 | 3 |
| STEM company founder | 5 | 6 | 0 | 0 | 5 | 3 |
| Computer or data scientist | 1 | 1 | 3 | 3 | 4 | 2 |
| Health or medical staff | 1 | 1 | 3 | 3 | 4 | 2 |
| Software developer | 3 | 3 | 1 | 1 | 4 | 2 |
| Faculty (within a STEM department) | 2 | 2 | 2 | 2 | 4 | 2 |
| Pharmacist | 2 | 2 | 1 | 1 | 3 | 2 |
| Research associate | 3 | 3 | 0 | 0 | 3 | 2 |
| Safety/security specialist<br>(within a STEM department) | 3 | 3 | 0 | 0 | 3 | 2 |
| Bioengineer | 1 | 1 | 1 | 1 | 2 | 1 |
| Technician | 0 | 0 | 2 | 2 | 2 | 1 |
| Mathematician | 0 | 0 | 1 | 1 | 1 | 0.6 |
| Nuclear scientist | 0 | 0 | 1 | 1 | 1 | 0.6 |
| Physical therapist | 0 | 0 | 1 | 1 | 1 | 0.6 |
| Dentist | 0 | 0 | 1 | 1 | 1 | 0.6 |
| Other <sup>b</sup> | 5 | 6 | 13 | 14 | 18 | 10 |

*Note:* The job held by one individual may be described by more than one category. <sup>a</sup> This category includes those individuals who are currently attending a research-based graduate program and work as research assistants. <sup>b</sup> In order to protect the identities of study participants, any job or professional activity held by SULI or CCI alumni outside of the STEM or health career pathways will not be described outside of the “other” category. This information is current through December 2021, which is 5 to 12 years after alumni participation in the CCI and SULI programs.

**Table S3.**
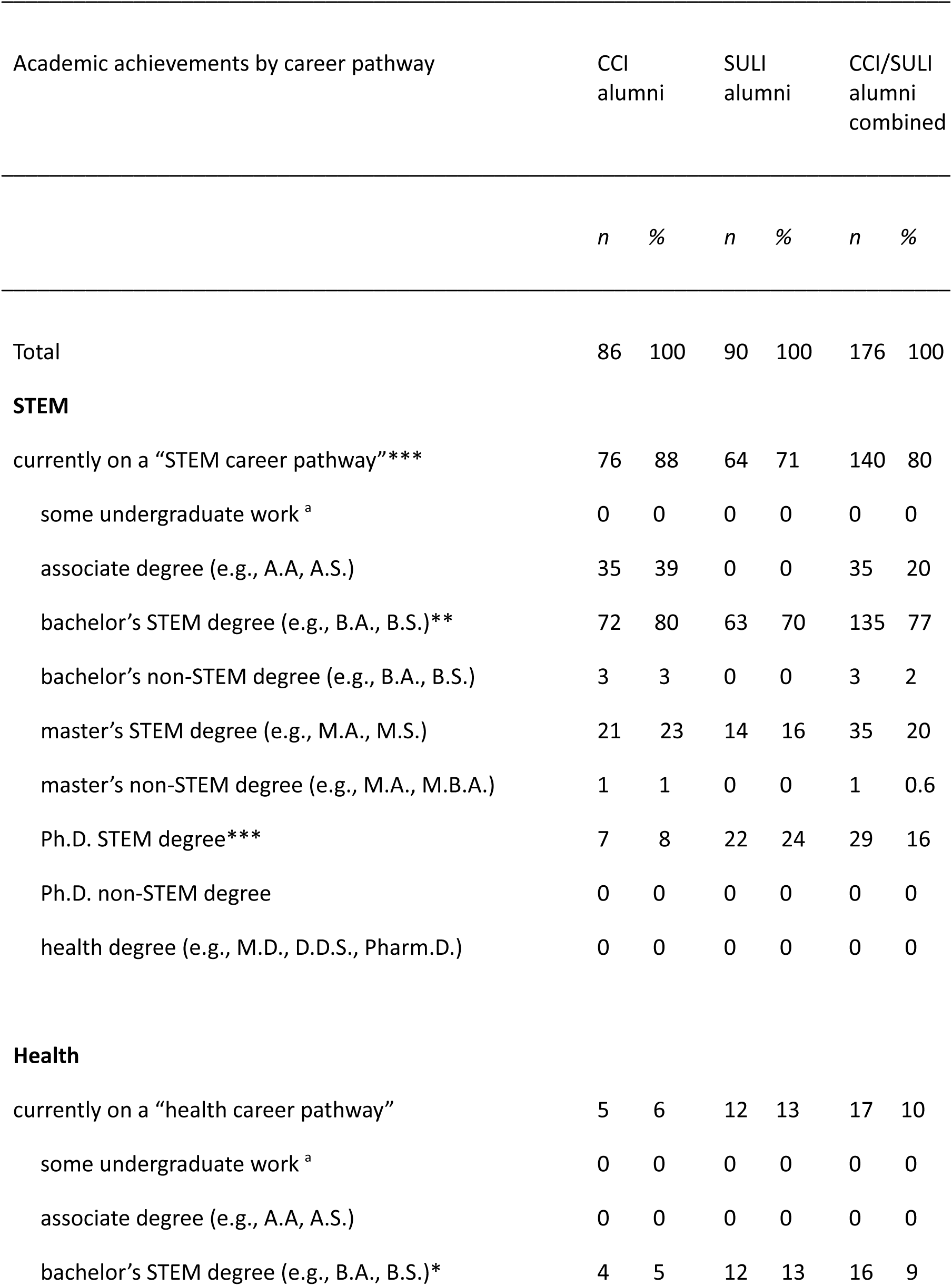
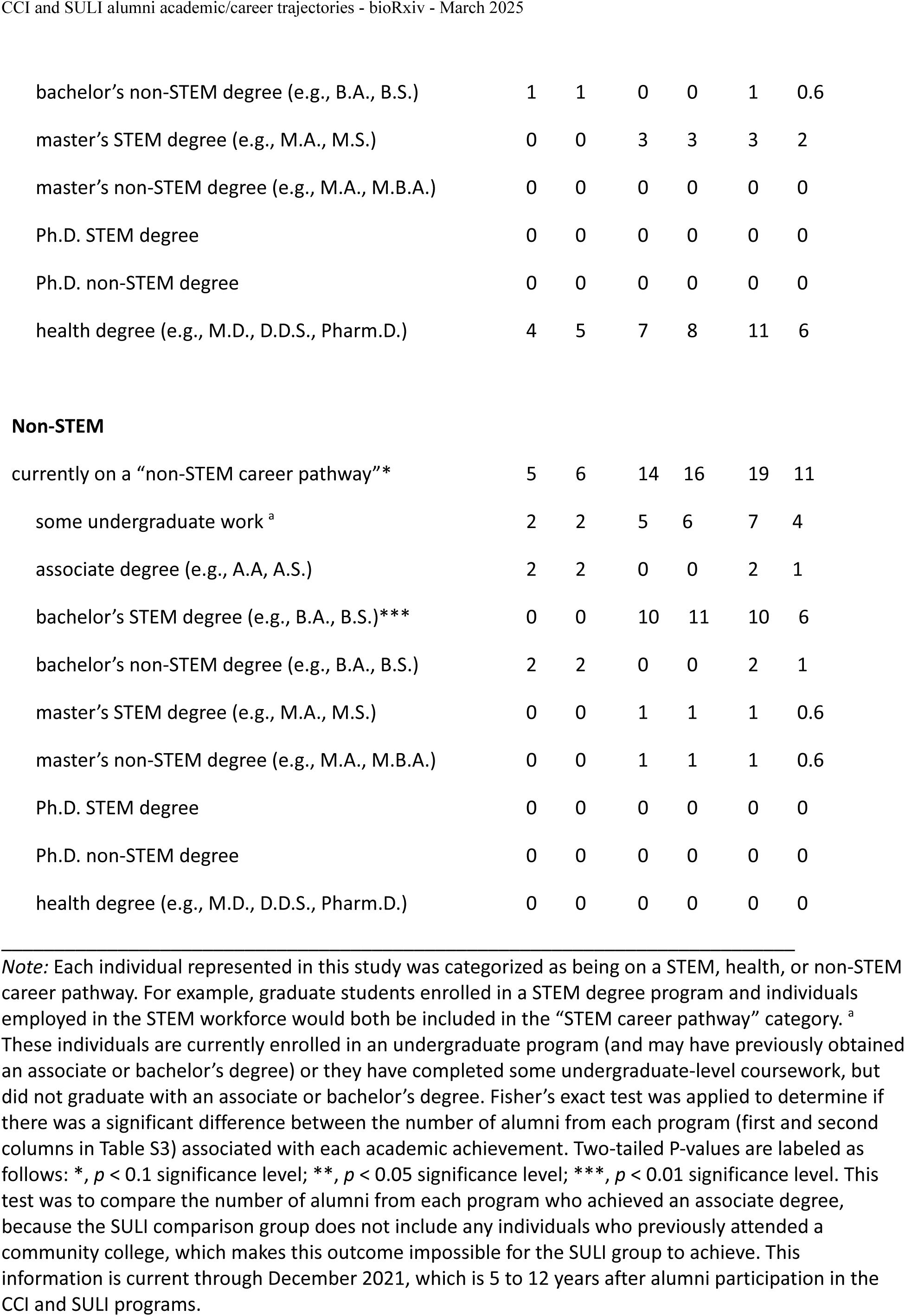
Academic Achievements of Study Participants by Career Pathway.

| Academic achievements by career pathway | CCI alumni |  | SULI alumni |  | CCI/SULI alumni combined |  |
| --- | --- | --- | --- | --- | --- | --- |
|  | *n* | % | *n* | % | *n* | % |
| Total | 86 | 100 | 90 | 100 | 176 | 100 |
| **STEM** |  |  |  |  |  |  |
| currently on a “STEM career pathway”*** | 76 | 88 | 64 | 71 | 140 | 80 |
| some undergraduate work <sup>a</sup> | 0 | 0 | 0 | 0 | 0 | 0 |
| associate degree (e.g., A.A, A.S.) | 35 | 39 | 0 | 0 | 35 | 20 |
| bachelor’s STEM degree (e.g., B.A., B.S.)** | 72 | 80 | 63 | 70 | 135 | 77 |
| bachelor’s non-STEM degree (e.g., B.A., B.S.) | 3 | 3 | 0 | 0 | 3 | 2 |
| master’s STEM degree (e.g., M.A., M.S.) | 21 | 23 | 14 | 16 | 35 | 20 |
| master’s non-STEM degree (e.g., M.A., M.B.A.) | 1 | 1 | 0 | 0 | 1 | 0.6 |
| Ph.D. STEM degree*** | 7 | 8 | 22 | 24 | 29 | 16 |
| Ph.D. non-STEM degree | 0 | 0 | 0 | 0 | 0 | 0 |
| health degree (e.g., M.D., D.D.S., Pharm.D.) | 0 | 0 | 0 | 0 | 0 | 0 |
| **Health** |  |  |  |  |  |  |
| currently on a “health career pathway” | 5 | 6 | 12 | 13 | 17 | 10 |
| some undergraduate work <sup>a</sup> | 0 | 0 | 0 | 0 | 0 | 0 |
| associate degree (e.g., A.A, A.S.) | 0 | 0 | 0 | 0 | 0 | 0 |
| bachelor’s STEM degree (e.g., B.A., B.S.)* | 4 | 5 | 12 | 13 | 16 | 9 |
| bachelor's non-STEM degree (e.g., B.A., B.S.) | 1 | 1 | 0 | 0 | 1 | 0.6 |
| master's STEM degree (e.g., M.A., M.S.) | 0 | 0 | 3 | 3 | 3 | 2 |
| master's non-STEM degree (e.g., M.A., M.B.A.) | 0 | 0 | 0 | 0 | 0 | 0 |
| Ph.D. STEM degree | 0 | 0 | 0 | 0 | 0 | 0 |
| Ph.D. non-STEM degree | 0 | 0 | 0 | 0 | 0 | 0 |
| health degree (e.g., M.D., D.D.S., Pharm.D.) | 4 | 5 | 7 | 8 | 11 | 6 |
| Non-STEM |  |  |  |  |  |  |
| currently on a "non-STEM career pathway"* | 5 | 6 | 14 | 16 | 19 | 11 |
| some undergraduate work <sup>a</sup> | 2 | 2 | 5 | 6 | 7 | 4 |
| associate degree (e.g., A.A, A.S.) | 2 | 2 | 0 | 0 | 2 | 1 |
| bachelor's STEM degree (e.g., B.A., B.S.)*** | 0 | 0 | 10 | 11 | 10 | 6 |
| bachelor's non-STEM degree (e.g., B.A., B.S.) | 2 | 2 | 0 | 0 | 2 | 1 |
| master's STEM degree (e.g., M.A., M.S.) | 0 | 0 | 1 | 1 | 1 | 0.6 |
| master's non-STEM degree (e.g., M.A., M.B.A.) | 0 | 0 | 1 | 1 | 1 | 0.6 |
| Ph.D. STEM degree | 0 | 0 | 0 | 0 | 0 | 0 |
| Ph.D. non-STEM degree | 0 | 0 | 0 | 0 | 0 | 0 |
| health degree (e.g., M.D., D.D.S., Pharm.D.) | 0 | 0 | 0 | 0 | 0 | 0 |

**Figure S1.**
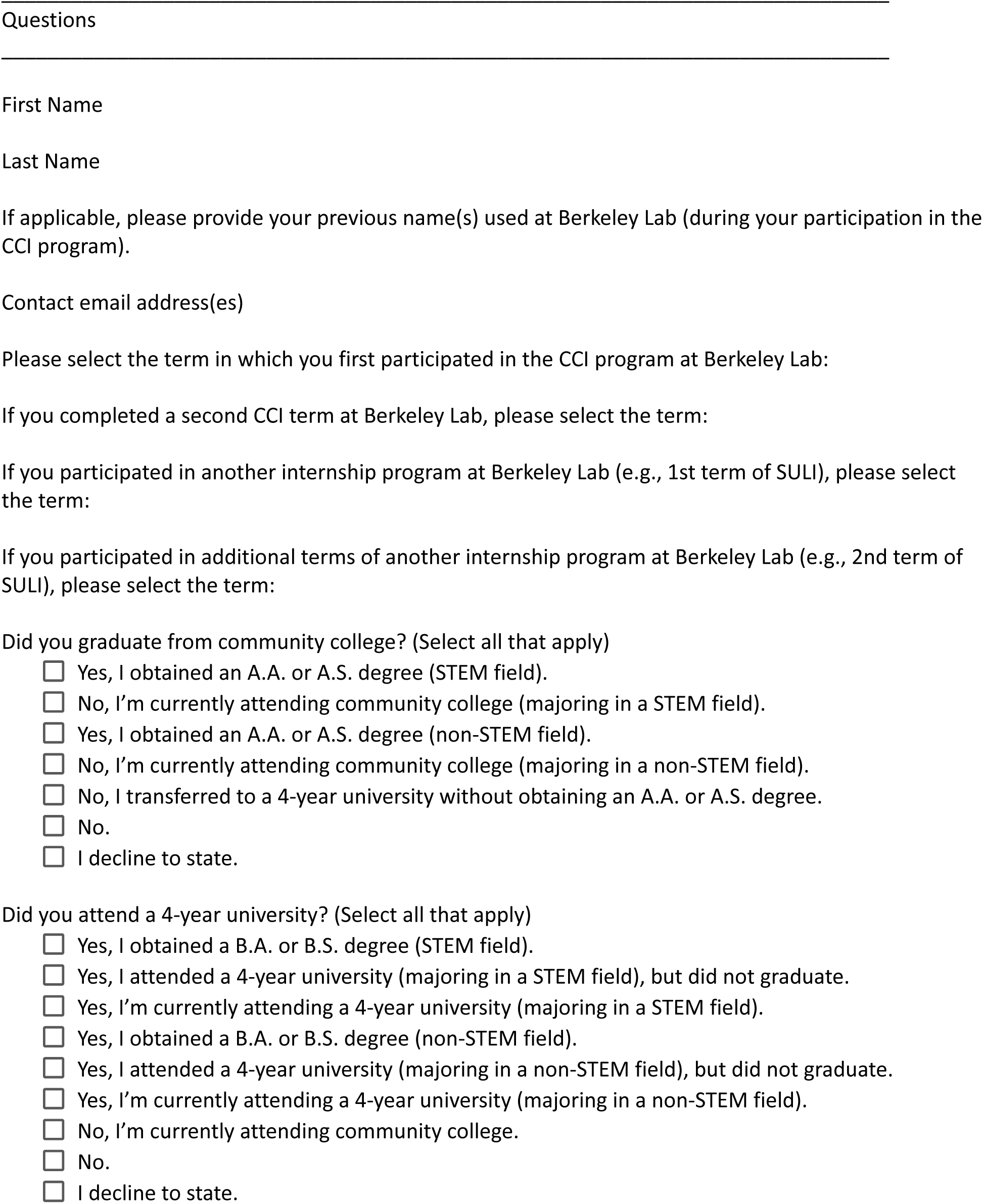
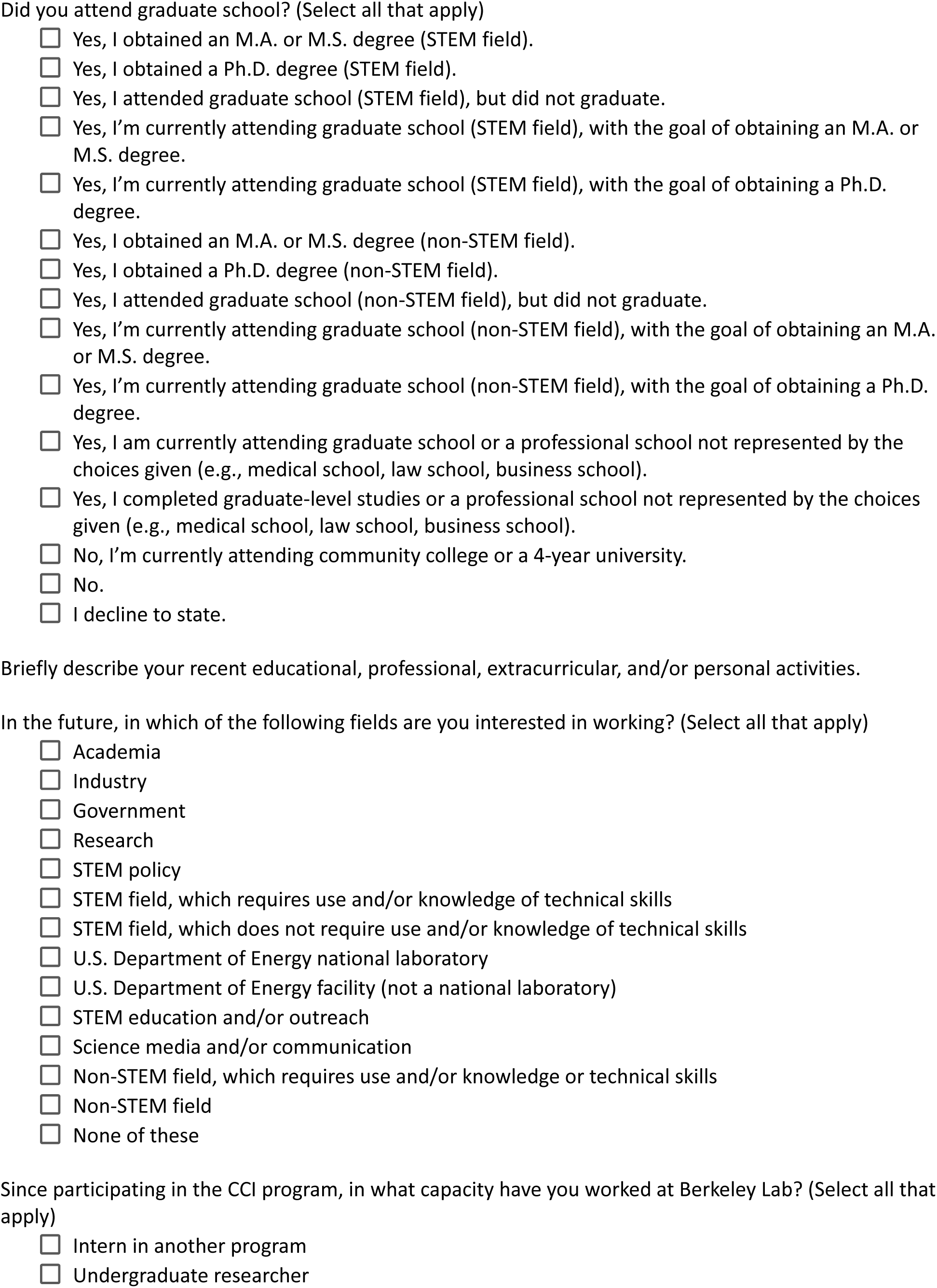
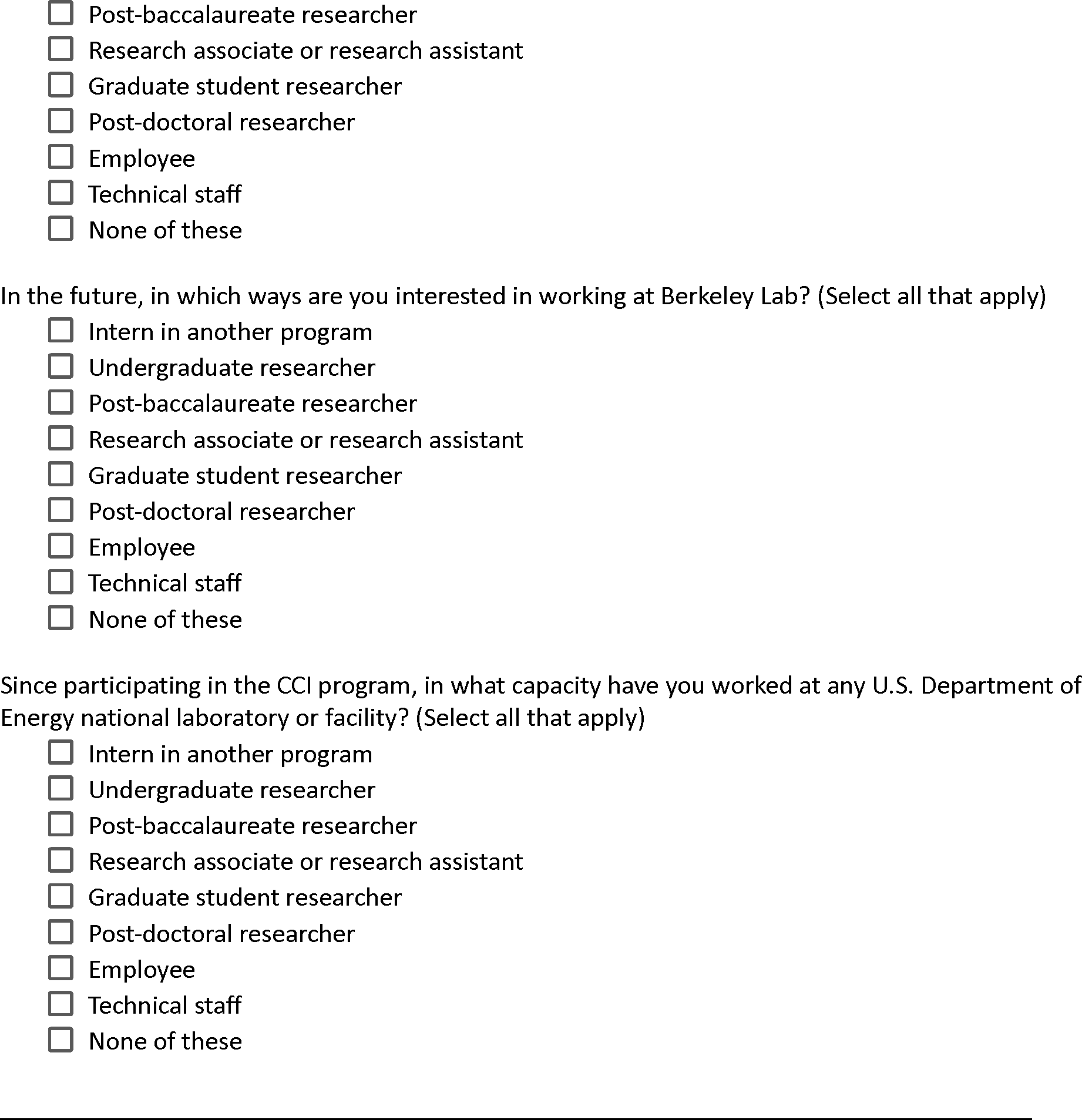
Selection of Questions from the Community College Internship (CCI) Alumni Survey.

**Figure S2.**
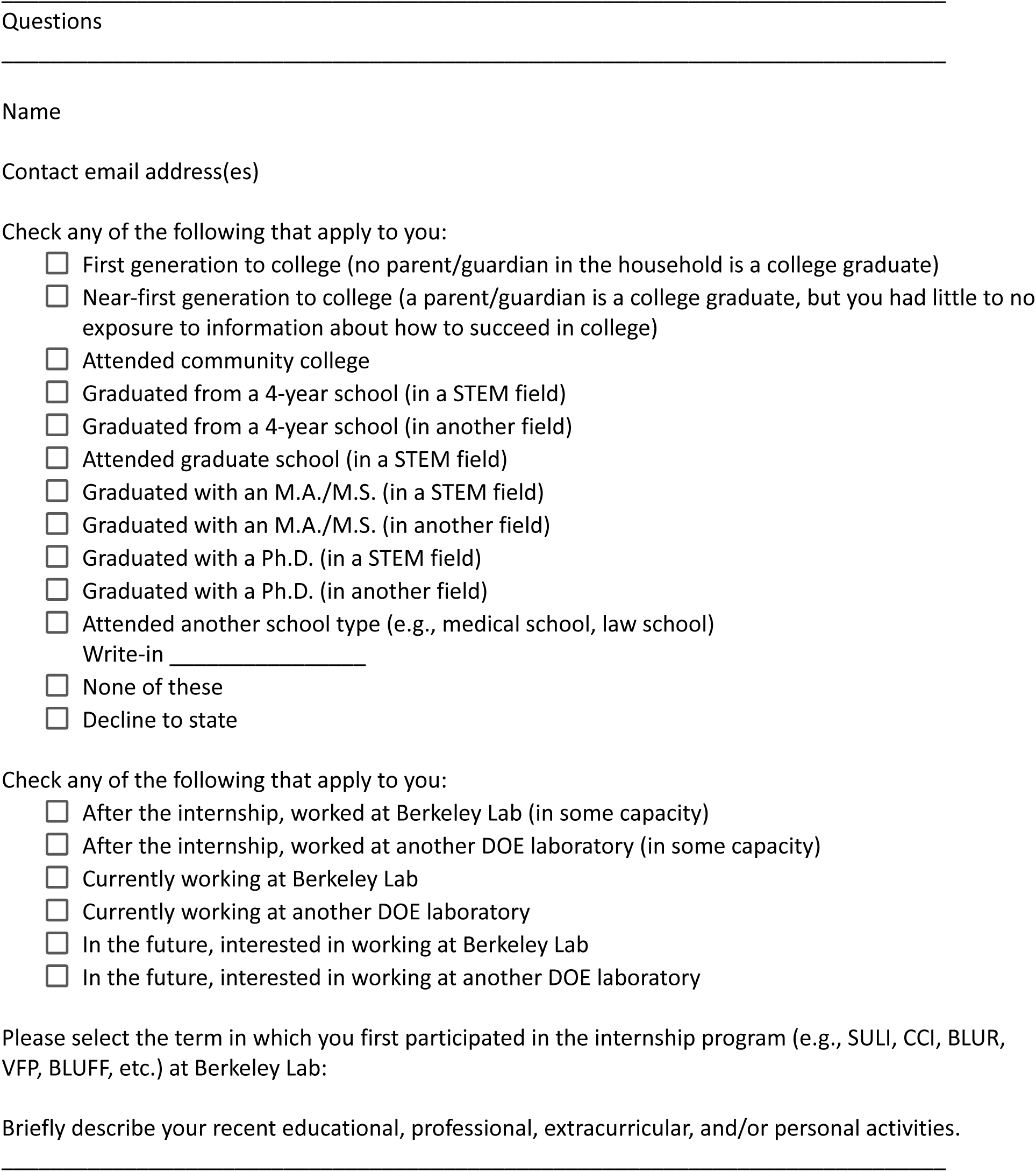
Selection of Questions from the <u>LBNL Internship Alumni Survey</u>.

**Figure S3.**
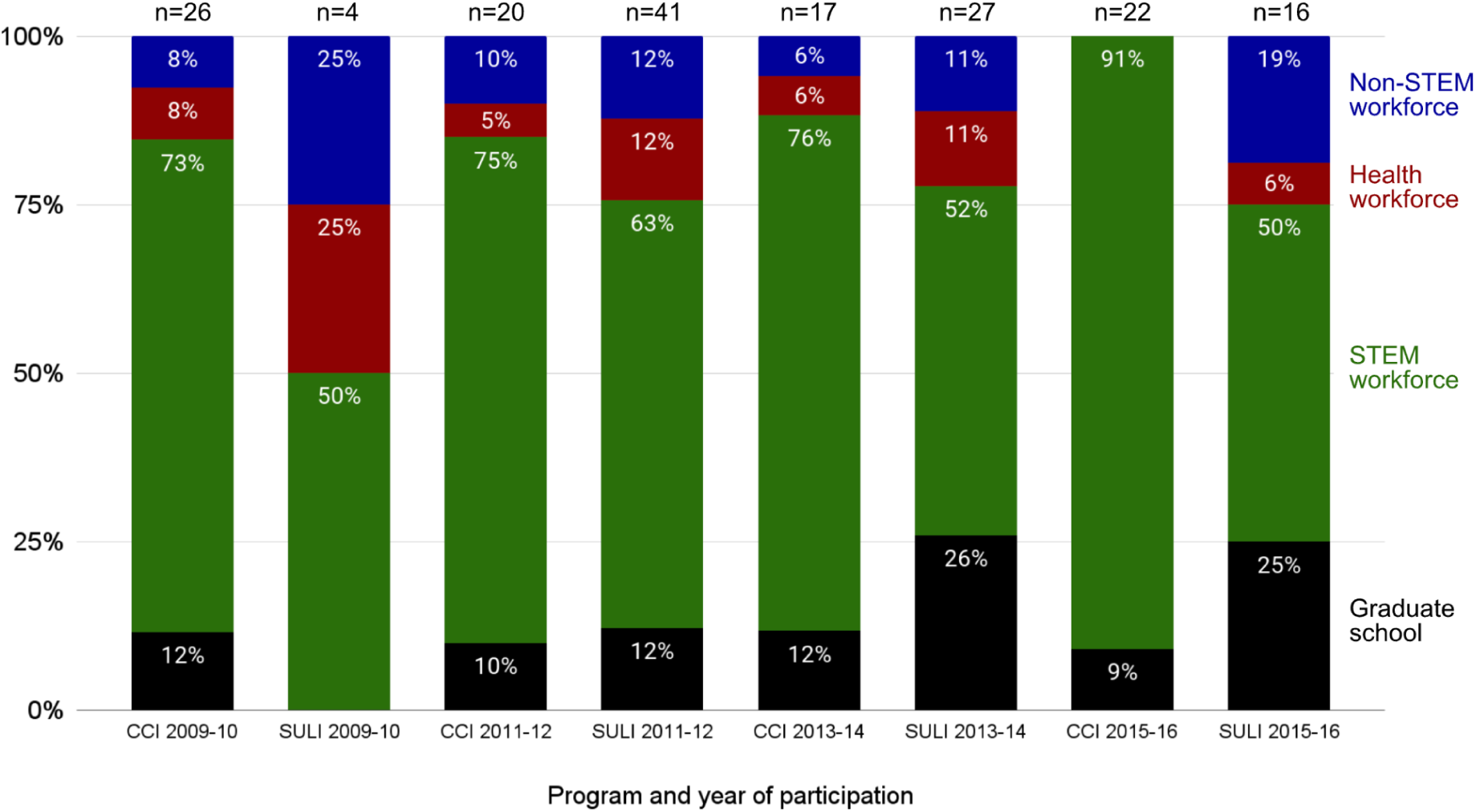
Current Academic and Career Activity for Study Participants by Time Frame and Program. *Note.* The graph shows which percentage of the CCI and SULI participants from a 2-year time frame are currently attending graduate school (any program type), or employed in the STEM, health, or non-STEM workforces. The time frames included are 2009-2010, 2011-2012, 2013-2014, and 2015-2016. This information is current through December 2021, which is 5 to 12 years after alumni participation in the CCI and SULI programs.

## Footnotes

1 We use the term “community college” to distinguish these higher education institutions that offer associate degrees, workforce education, and certificate programs from those that offer bachelor’s and graduate degrees (referred to as “baccalaureate granting institutions”). However, the U.S. has various institution types that fall into this category, including city colleges, community colleges, junior colleges, technical colleges, technical schools, trade schools, vocational schools, etc.

2 We avoid referring to baccalaureate granting institutions as “4-year” institutions or community colleges as “2-year” institutions, because undergraduates spend varying amounts of time at these institutions in pursuit of their academic goals (Cohen & Kelly, 2019; Complete College America, 2014; Jain et al., 2020; Ocean et al., 2022).

3 In this study we are using the Method Reporting with Initials for Transparency (MeRIT) approach as described by Nakagawa and colleagues (2023). A.M. = Aparna Manocha, A.M.B. = Anne M. Baranger, A.N.Z. = Astrid N. Zamora, E.W.L. = Esther W. Law, G.O.M. = Gabe Otero Munoz, J.J.S. = Julio Jaramillo Salcido, L.E.C. = Laleh Coté, and S.V.D. = Seth Van Doren.

## Notes

### Competing Interest Statement

The authors have declared no competing interest.

